# ATLS2021-PA (A Sphingolipid mimetic): Upcoming Host Directed Therapy for Acute and MDR Tuberculosis

**DOI:** 10.1101/2025.03.17.643690

**Authors:** Niharika Sharma, Rahul Sharma, Vijay Hadda, Amit Kumar Singh, Hridayesh Prakash

## Abstract

Several reports have shown that reduced Sphingolipids content in the lungs largely contributes to the outcome of Tuberculosis diseases. In this context, our pioneer study has advocated that Sphingosine −1 phosphate (a central Sphingolipid metabolite) can help host in limiting *Mycobacterium tuberculosis* burden in lungs by their virtue of tweaking M1 retuning of infected macrophages. This indicated that S1P may serve as anti-tubercular regimen, however allergic and autoimmune manifestation of S-1P refrain its further use as anti-tubercular Drugs. In view of this limitation, and high demand of developing newer host directed anti tubercular regimen, we explored whether enhancing / boosting Sphingolipids levels via *de novo* pathways by our patented drugs ATLS2021 and PA would also help host in controlling mycobacterial infection. Our results prudently demonstrated that ATLS2021 was effective in controlling survival of both wild type and MDR strain of *TB* in various models we tested. Our results further demonstrated the influence of ATLS2021 on NO mediated killing of mycobacteria which indicated one of most possible anti-mycobacterial mechanism of ATLS2021. Additionally, ATLS2021 lowered the IC50 value of Rifampicin for both wild-type and MDR TB by sensitizing the MDR clinical isolate of *Mycobacterium tuberculosis* to Rifampicin-mediated killing. ATLS2021 induced immunogenic responses in blood derived CD14+ macrophages from both healthy donors as well as MDR-TB patient demonstrating immune adjuvant potential of ATLS2021. Real time PCR data demonstrated that ATLS2021 enhanced the expression of almost all key enzymes involved in the Sphingolipid biosynthesis pathways in the CD14 positive monocytes from healthy donors and MDR TB patients. Taken together these results potentially advocated that ATLS2021 based approach is potentially immunogenic interventions against TB.

## Introduction

Immunocompetent host limit the mycobacterial infection in granulomatic lesions which is an indicative of host immunity against infection. This depends on the gradient of Sphingolipids which are amphipathic and dual specific lipids which can favour both host and pathogen (1, 2) during the acute / latent infection (3). Several studies (4) including ours have demonstrated the active involvement of Sphingolipids on host innate response against *Mycobacterial infection* (5) 28400772. Of particular note our pioneer and compelling study (6) has advocated that Sphingosine −1 phosphate; the key Sphingolipid moiety is capable of controlling *M. tb* infection. Since metabolic programming of host limits effective control of variety of infections including *M. tb*. Among various micronutrients, host Sphingolipid play a deceive role in the pathophysiology of respiratory diseases including COPD, asthma and Interstitial lung diseases. Several reports have shown that Sphingolipids content in Tuberculosis (TB) patients reduces (7) with disease progression (8) because host utilize Sphingolipids for the management of the disease while maintaining homeostasis. As a result, TB patients become poor in their Sphingolipids metabolites like S-1P which is anti-tubercular in nature. Hence low gradient of pulmonary and serum Sphingolipid index confer poor immunity against mycobacteria and promote diseases severity (9). On account of this, we argued whether supplementing host with S-1P would enhance innate immunity and control the infection. Supporting this argument, our pioneer study have prudently demonstrated therapeutic impact of S1P in controlling *M. tb* burden in host (6) warranting its application as host directed novel anti tubercular regimen. However, due to several acute inflammatory and allergic manifestations which are associated with pure S-1P, in spite of its anti-tubercular nature (10) it may not qualify the said criteria for clinics.

In view of high surge of developing newer host directed anti tubercular regimen, development of an alternative approach for enhancing levels of Sphingolipids production (11) is paramount for affording protection against Mycobacterial diseases which are increasing exponentially. In this regard, we have adopted ATLS2021 based *de novo* approaches for enhancing Sphingolipids synthesis in host for clearing infection efficiently. Following our expectations, our study has prudently demonstrated strong anti-mycobacterial influence of ATLS2021 (precursor of Sphingolipids *de novo* synthesis) against various strain of mycobacteria we used including clinical isolate of MDR-TB. This indicated that ATLS2021 based approach is certainly efficient strategy for boosting immunity against infection. Moreover, ATLS2021 has sensitized both wild type and clinical isolate of MDR tuberculosis for 1^st^ and 2^nd^ line anti tubercular drugs. Most intriguingly, ATLS2021, while controlling Mycobacterial growth in both broth cultures (cell culture system) as well as *in vivo*or *ex vivo*?, also mitigated infection induced inflammatory response suggesting that ATLS2021 may qualify pharmacological criteria for being used either as pro-drug / adjunct to the existing ATD. In line with these result, animal data also demonstrated that ATLS2021 alone or in combination of PA was able to reduce Mycobacetrial burden significantly in comparism of infection control. As expected, while ATLS2021 supplementation enhanced pulomanry titre of TNF-α, it concomitantly inhibited IL-10 titre in the lungs of infected animals indicating immunogenic programming potential of ATLS2021 in the infected animals. Our prospective clinical study with further shed light on the immunogenic potential of ATLS2021 on blood derived CD14+ macrophages from both healthy donors as well as MDR-TB patient. ATLS2021 enhanced generation of NO on its own but also augmented LPS and Rifampicin induced NO levels in the culture supernatant of CD14+ Monocytes. This observation was further substantiated by IFN gamma titre which was also augmented by ATLS2021. Most interestingly ATLS2021 supplementation inhibited the secretion of both IL-10 & IL-6 titre by the macrophages from MDR patients indicating immune conditioning potential of ATLS2021 in blood derived CD14+ monocytes which are involved in the protection against TB. Real time PCR data further demonstrated that ATLS2021 enhanced the expression of almost all key enzymes involved in the Oxy-Sphingolipid biosynthesis pathways in the CD14+ monocytes from healthy donors and MDR TB patients. Taken together these results potentially advocated that ATLS2021 based approach is potentially immunogenic Interventions against mycobacterial diseases. We believe that our approach is transnationally viable and would add significantly to the existing host directed therapies against drug resistant mycobacterial infection.

## Results

### ATLS2021PA kills mycobacteria directly

To access the anti-mycobacterial influence of ATLS2021, we first analyzed the impact of ATLS2021 on the growth of *Mycobacterium smegmatis (M. smegmatis)*. For this purpose 1×10^6^ *M. smegmatis* were grown in 7H9 broth media (containing Glycerol) in bacterial shaker incubator shaker fixed at 37 ^°^C and 120 RPM in presence or absence or various concentration of ATLS2021 for 24^th^ and 48^th^ **(Figure 1a)** intervals. Following our expectation, ATLS2021 was able to restrict the growth pattern of *M. smegmatis* at higher dose by more than 50% of sham respectively at both at 24 & 48hr intervals. These data demonstrated that ATLS2021 is effective in controlling *M. smegmatis*. To determine whether addition to PA would restrict the growth of *M. smegmatis* further, *M. smegmatis* cultures were grown in presence of ATLS2021 along with PA at indicated concentration. Following our prediction, addition of PA influenced anti mycobacterial impact of ATLS2021 further **(Figure 1b)** and unambiguously demonstrated that ATLS2021 alone is effective in controlling *M. smegmatis* growth.

**Figure 1:**
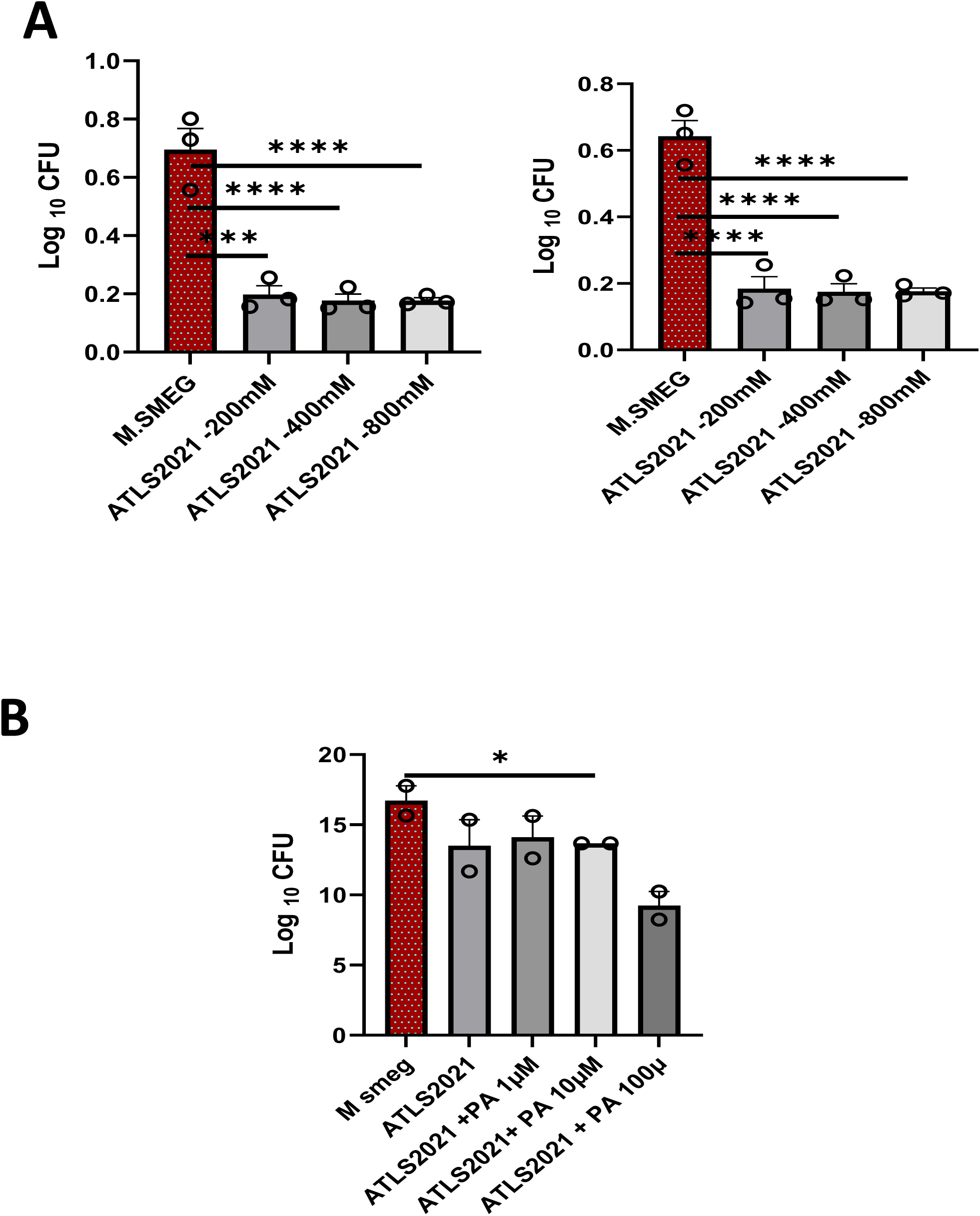
ATLS2021-PA kills mycobacteria directly. M smegmatis cultured were grown in presence of ATLS2021+PA and their CFU was quantified by OD based methods at 24h **(A)** and 48 hours **(B)**. Shown here the mean of CFU ± SEM quantified from 4 independent repeats at indicated time intervals.

### ATLS2021PA induces RNI and Hypoxia for killing of mycobacteria

Macrophages are the main reservoir of mycobacterial pathogen in host and utilize both ROS and RNI (full form?) for killing Mycobacteria in their phagolysosomal compartment. Above results clearly indicated anti-mycobacterial role of ATLS2021 and raised a possibility that ATLS2021 may utilize these arms for killing mycobacteria. To this end, we first analysed the impact on ATLS2021 on NO dependent killing of mycobacteria. To that purpose, we expanded *M. smegmatis* cultures in presence of Sodium Nitroprusside (SNP) (NO donor). As per our expectation, SNP alone inhibited the growth of *M. smegmatis* **(Suppl Figure 1)** and most interestingly, supplementation of SNP treated *M. smegmatis* cultures with ATLS2021 further reduced the growth of *M. smegmatis* exponentially **(Figure 2a)** which revealed possible influence of ATLS2021 and NO mediated killing of mycobacteria. Other than RNI, cellular hypoxia is important for the post killing clearance of mycobacterial clearance for homeostasis. Since ATLS2021 is potentially homeostatic in nature so we presumed that ATLS2021 will enhance / mimic in line with hypoxia and promote mycobacterial killing further. To demonstrate this, we expanded mycobacterial cultures in presence of CoCl_2_ which stabilized HIF1 and manifest hypoxia in cells. CoCl_2_ inhibited mycobacterial growth significantly **(Suppl Figure 2b)** supporting our hypothesis. Most interestingly, addition of ATLS2021 further inhibited the growth of mycobacteria in CoCl_2_ treated cultures **(Figure 2b)**. These findings clearly revealed that ATLS2021 promote both redox apparatus and may help in clearing the infection in host. This is very important aspect because infected macrophages, during clearing infection undergo phenotypic switch where hypoxia promote not only bacterial clearance but also resolution of infection induced inflammatory response indicating its impact of homeostatic mechanism post infection. Following this presumption, our results clearly indicated synergistic impact of SNP and CoCl_2_ in controlling mycobacterial growth **(Suppl Figure 3a).** Most Interestingly, these combination promoted the anti-tubercular action of Rifampicin **(Figure 2b & Suppl Figure 3b)**suggesting that these are most pertinent component of host which assist anti tubercular drugs for killing mycobacteria effectively.

**Figure 2:**
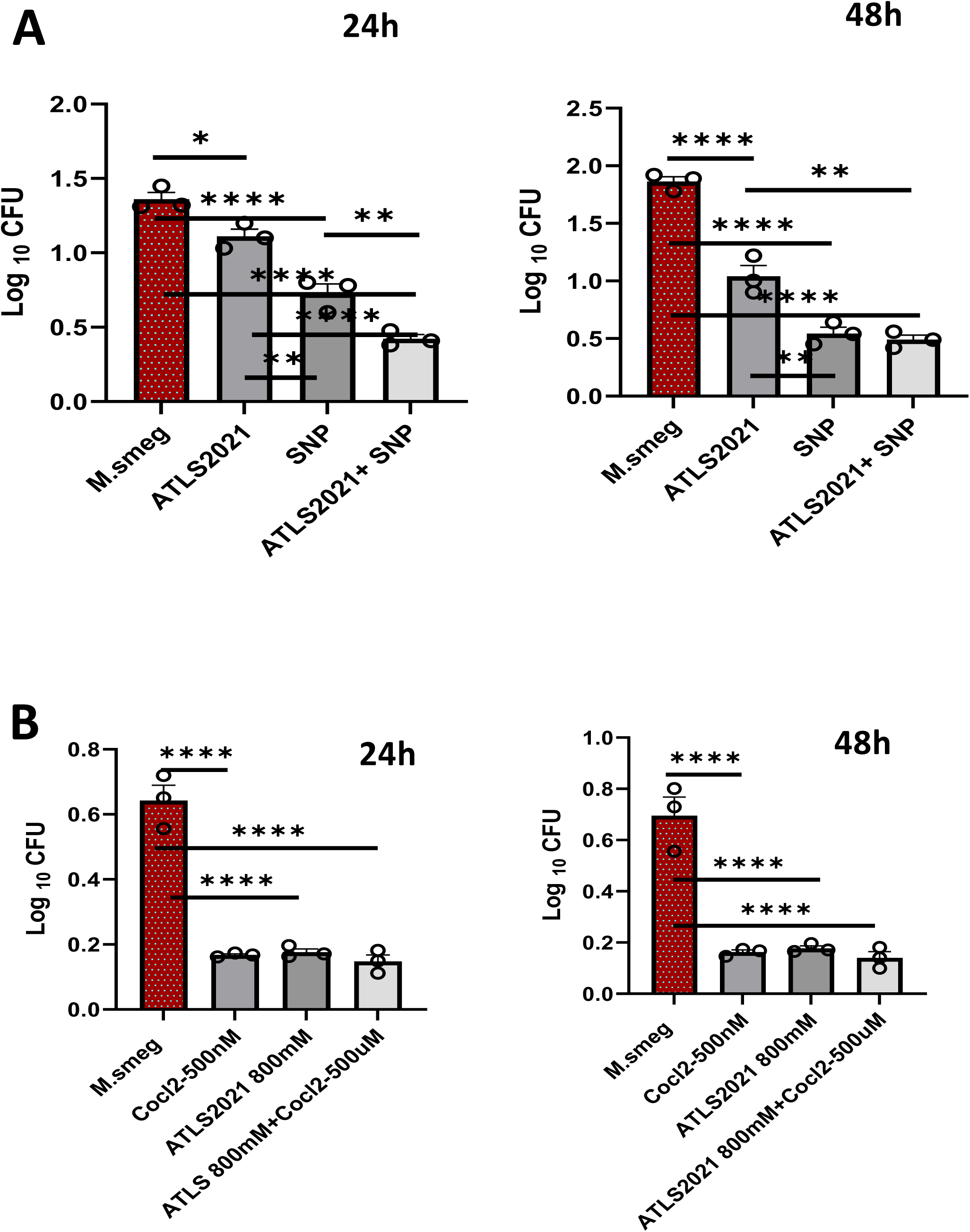
ATLS2021 mimic RNI and hypoxia for killing mycobacteria. ATLS2021 pulsed mycobacterial cultures were treated with NO donor (Sodium Nitroprusside) and Hypoxia mimetic CoCL2 and the impact of these stimulation were analysed on bacterial growth by quantifying their CFU at 24h **(A)** and 48 hours **(B)** which clearly indicates that ATLS2021 use RNI and Hypoxia for killing mycobacteria directly. Shown here the mean of CFU ± SEM quantified from 4 independent repeats at indicated time intervals.

### ATLS2021PA sensitizes mycobacteria for anti-tubercular drugs

On the basis of above results, we clarified whether ATLS2021 would alter the sensitivity of *M. smegmatis* sensitivity toward the anti-tubercular drugs or not. To demonstrate this, we cultured mycobacteria in presence of ATLS2021 with and without Anti Tubercular Drugs and assessed the growth pattern of mycobacteria. Following our prediction ATLS2021 enhanced anti tubercular potential of both Rifampicin and Isoniazid indicating adjunct potential of ATLS2021 **(Figure 3a & 3b).** These results provoked up to test further whether ATLS2021 can alter the sensitivity of various mycobacterial strains for Rifampicin or not. Following our expectations, ATLS2021 sensitized *M. smegmatis* for Rifampicin mediated killing indicating the adjunct nature of ATLS2021 and reduced the IC_50_ value of Rifampicin. On the basis of this, we further tested whether ATLS2021 would be effective against *M. tb* also. Supporting our presumption, ATLS2021 at 50mM concentration completely inhibited the growth of *M. tb* **(Figure 3c)** and to our good surprise to clinical isolate of MDR TB **(Figure 3d)** as well. These data potentially addressed the key question that ATLS2021 is effective not only against lab strain of Mycobacteria but also for clinical isolate of MDR TB.

**Figure 3:**
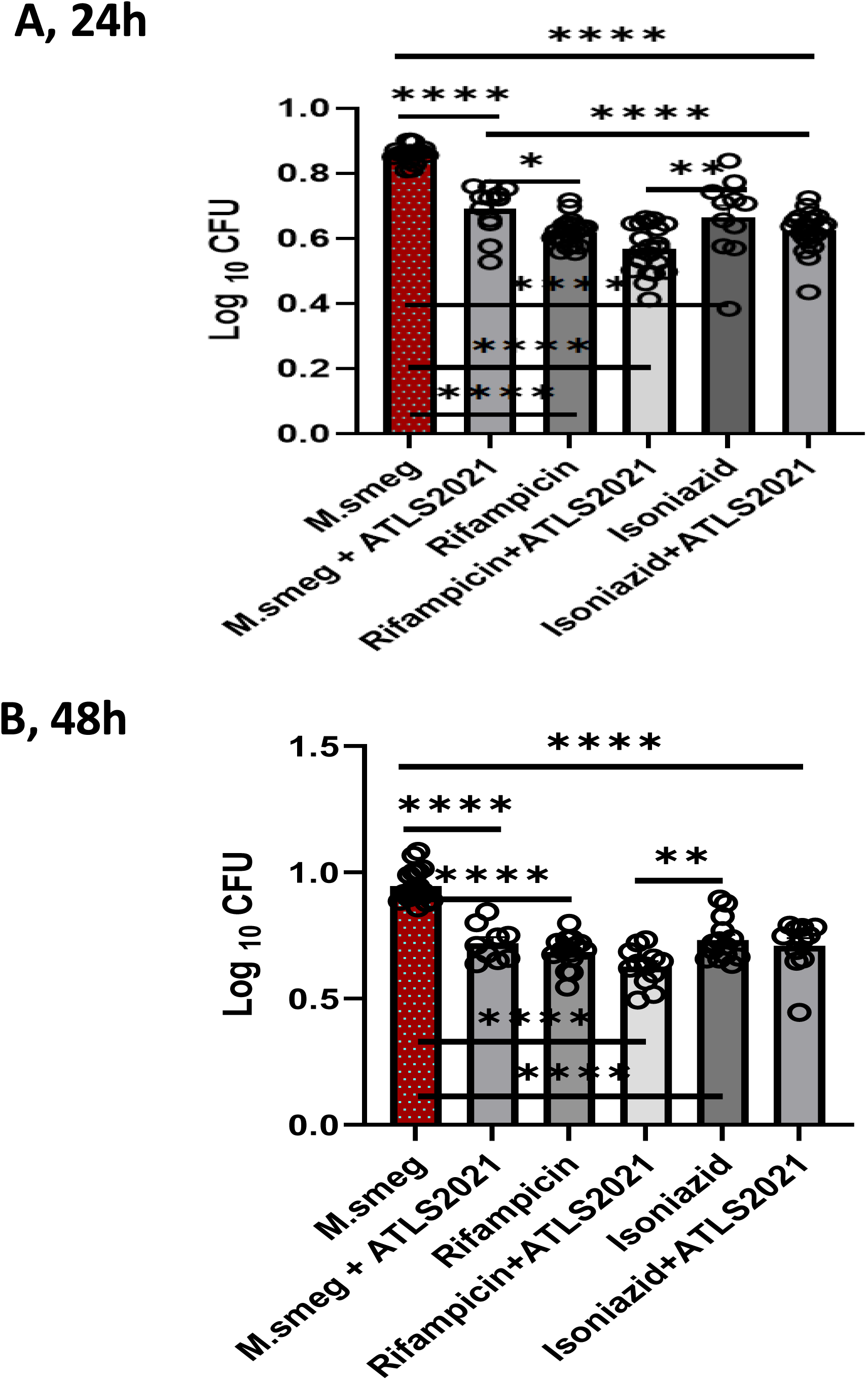
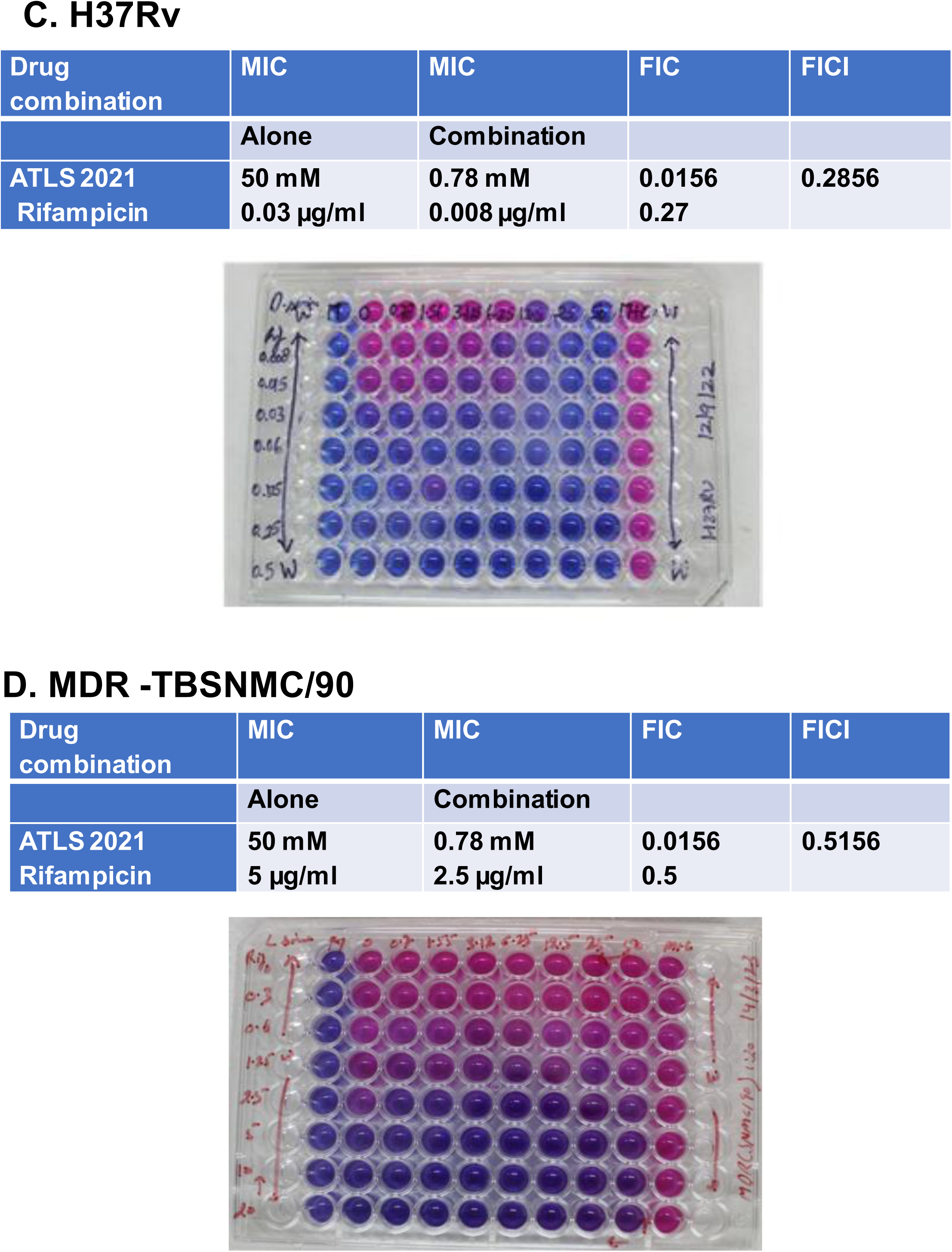
ATLS2021 is adjunct to Anti tubercular drugs. Mycobacterial cultures were pulsed with ATLS2021 in presence of anti-tubercular drugs and impacts of these stimulations were monitored on mycobacterial growth both at 24h (A) and 48h (B). CFU clearly indicates the adjunct potential of ATLS2021 on Rifampicin and Isoniazid. Out of both these ATD, ATLS2021 was more effective with Rifampicin. **(C)** In order to ensure one directional killing of Mycobacteria by this stimulation REMA method was followed as per the method mentioned in the text elsewhere. For accessing broad impact on ATLS2021 we used **M. tuberculosis**; H37Rv(C) and clinical isolate of **MDR TB; SNMC/90 (D)** MIC of RIF against all strains were calculated as per the scheme described in the text. Shown here is the representative result of 5 independent repeats.

### ATLS2021PA promotes killing of variety of Mycobacteria by variety of macrophages

Based on the above results, we expected that ATLS2021 would inhibit mycobacterial growth in macrophages as well. To demonstrate this, we used RAW 264.7, CD11b+/ Gr-1(−) peritoneal and Bone marrow derived macrophages, THP-1 and Blood derived CD14+ Monocytes populations. RAW264.7 macrophages were infected with *M. smegmatis* at MOI^−10^ in presence of ATLS2021 which was given in both prophylactically (one hour before) and therapeutically (4h post infection time intervals). Our initial set experiments, following our hypothesis, revealed that supplementation of exogenous ATLS2021 inhibited the growth of *M. smegmatis* replication not only in RAW264.7 **(Figure 4a)** but also in mouse peritoneal **(Figure 4b)** and Bone marrow derived macrophages **(Figure 4c)** also which is most probably due to increased synthesis of Oxy-Sphingolipid in ATLS2021 supplemented macrophages. However, ATLS2021 could not influence the mycobacterial uptake **(Suppl Figure 4)** by none of these macrophage’s populations. Interestingly addition of PA like before influenced the anti-mycobacterial potential of ATLS2021 against exponentially growing cultures of *M. smegmatis* **(Figure 5a)**. Interestingly both of them restricted the intracellular growth of *M. smegmatis* **(Figure 5b)** and supported microbiological data. Our further experiments with H37Rv further demonstrated that ATLS2021 & PA (100 µM) completely blocked the growth of H37Rv in RAW macrophages both at 48-& 72-hr post infection **(Figure 5c & 5d)** demonstrating their efficacy against pathogenic mycobacteria also. Interestingly at these concentration ATLS2021 and PA they cleared the information almost completely when administered as adjunct to antibiotic. Anti-mycobacterial impact of ATLS2021 in murine macrophages prompted us to rationalized that ATLS2021 would be effective in human monocyte / macrophages also. To demonstrate this, we infected human monocytic cells with H37Rv in presence of ATLS2021PA (confirm again)and anti-tubercular drugs. In line with RAW macrophages, CFU analysis of infected THP-1 also revealed adjunct potential of ATLS2021PA to anti-tubercular drug we tested. Although, ATLS2021 alone inhibited the growth of H37Rv in THP-1 only marginally however reduced this by 1.5 folds in combination of Anti tubercular drugs **(Figure 6a & 6b)** indicating its adjunct potential. On the basis of this, we raised question whether ATLS2021 would be effective in blood derived CD14+ macrophage or not. To assess this assess this we purified CD14+ monocytes and differentiated them with Human AB serum for five days and infected them with H37Rv in presence of ATLS2021 alone or in combination of Anti tubercular drug. Following our prediction and in line with RAW macrophages, ATLS2021 alone inhibited the growth of H37Rv and enhanced anti-tubercular potential of Anti tubercular drugs **(Figure 6c& 6d)** in CD14+ / CD16-human macrophages populations as well.

**Figure 4:**
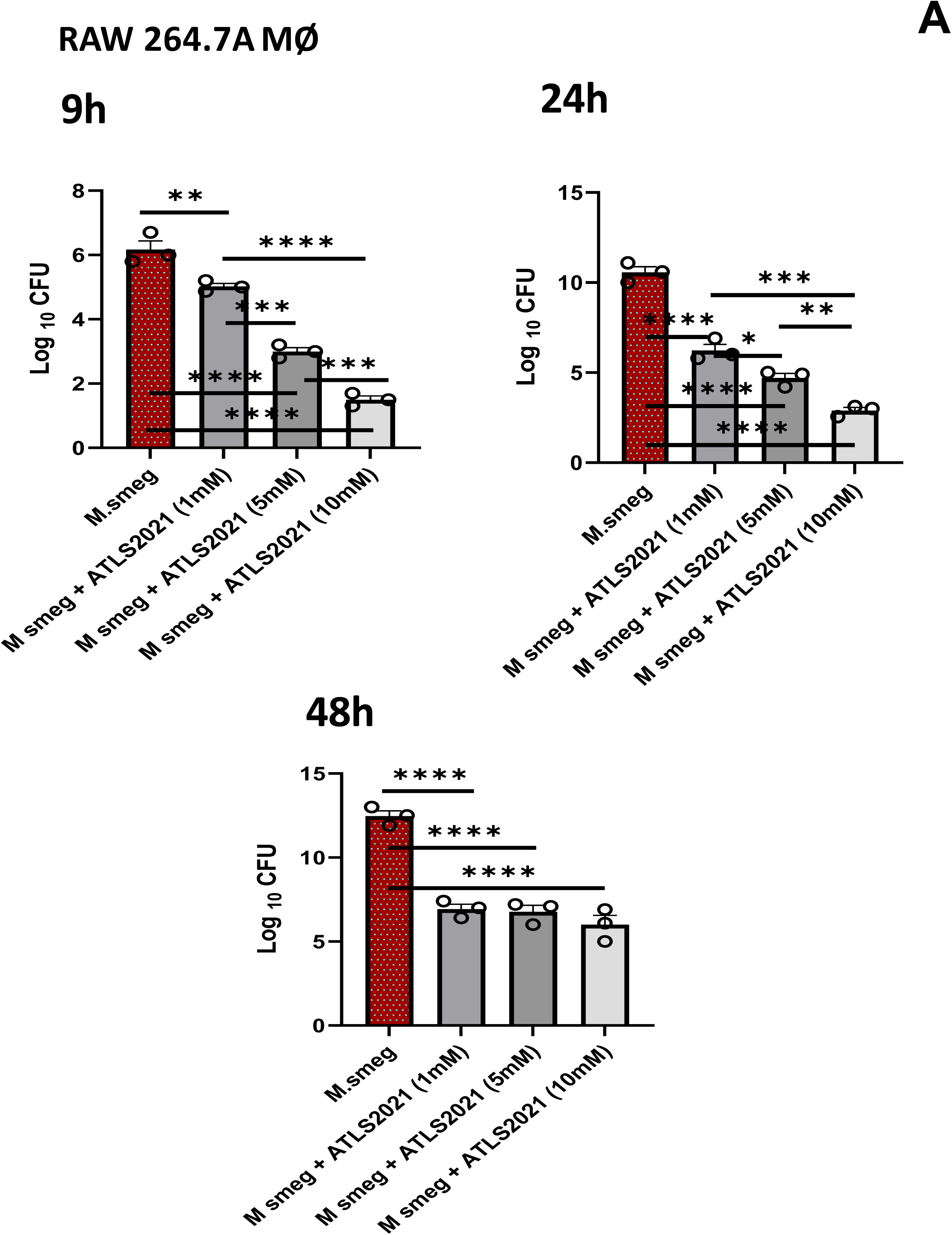
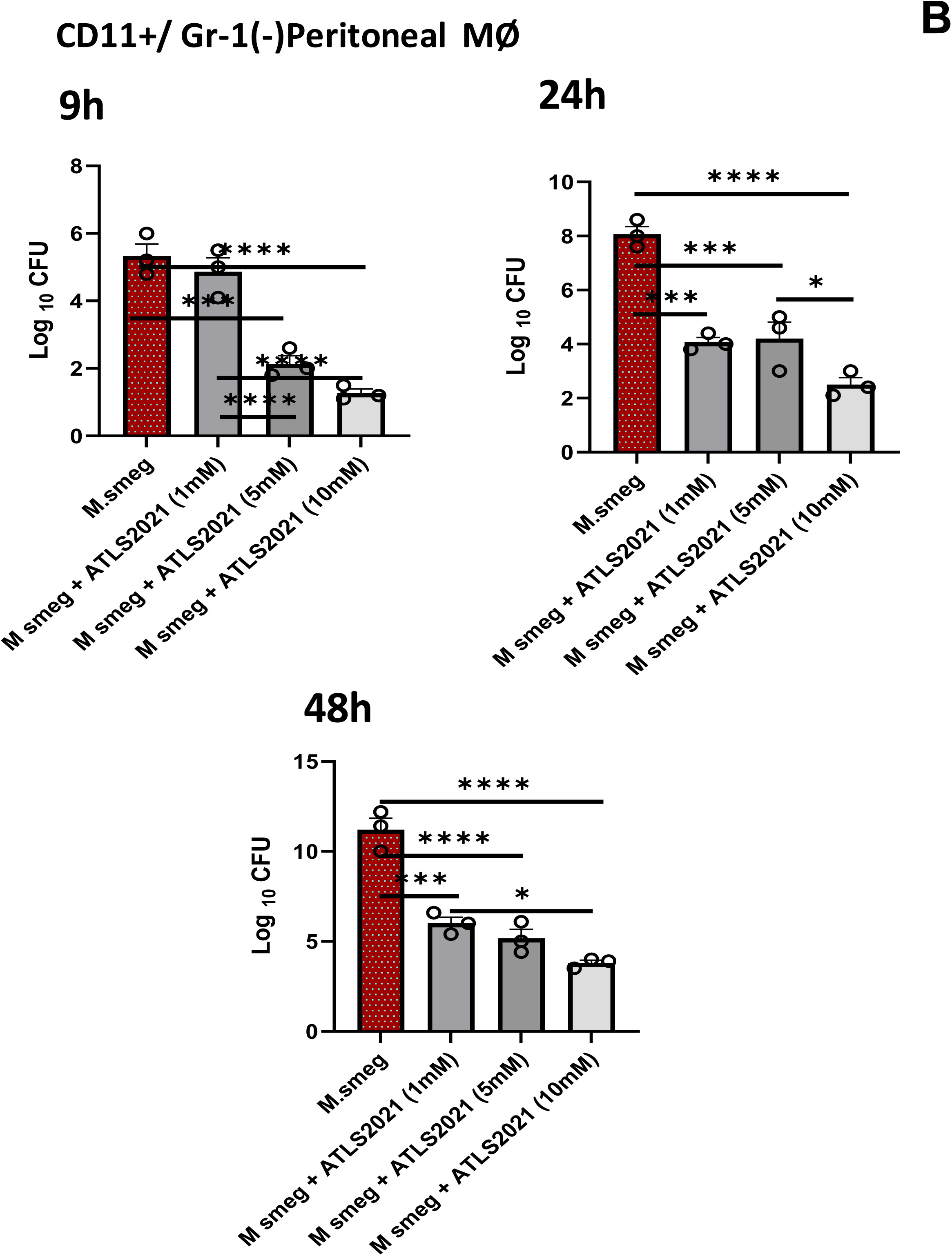
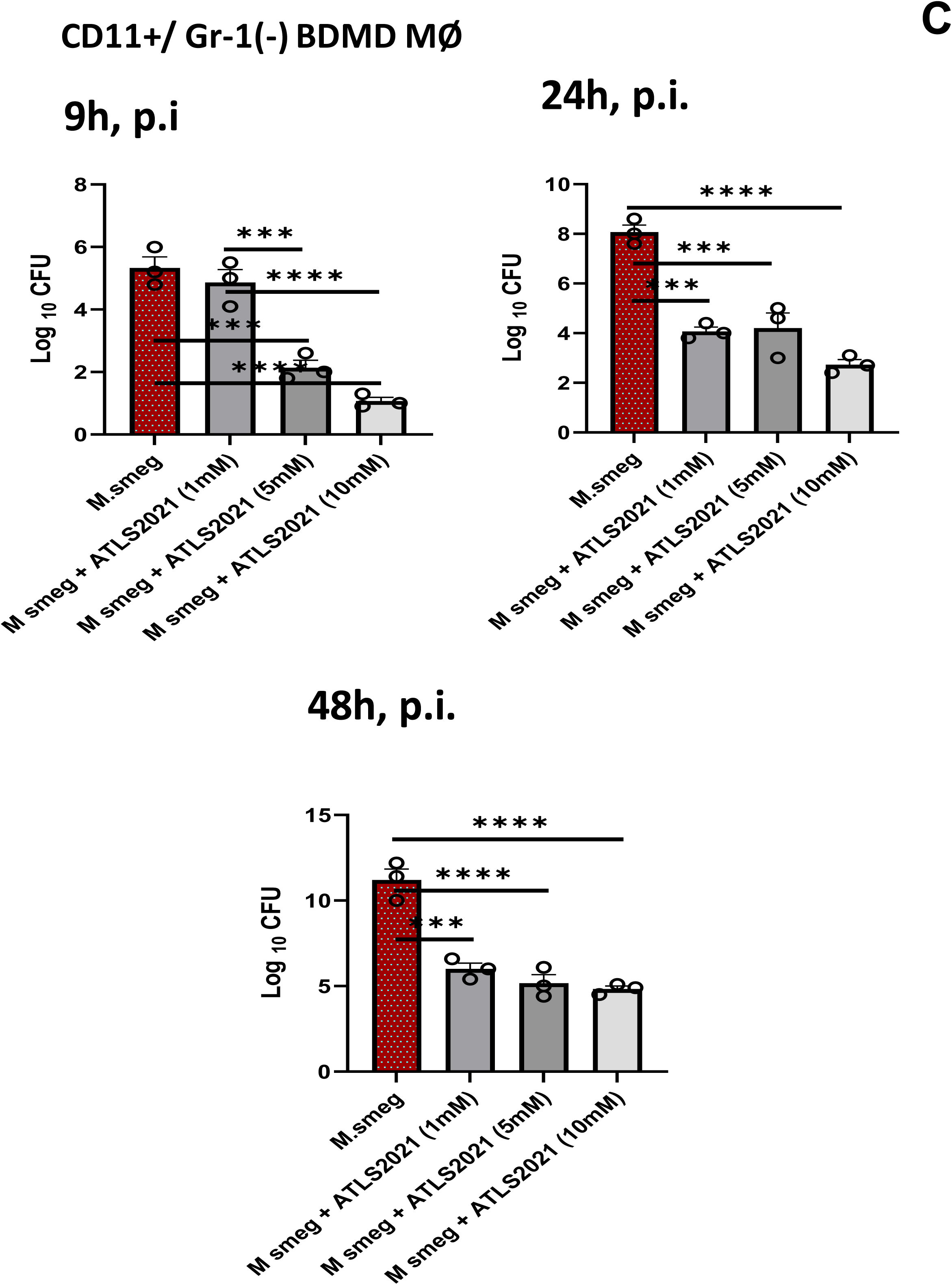
ATLS2021 control mycobacterial burden in vivo. Anti-tubercular potential of ATLS2021 was accessed in RAW 267.4A **(A)**, CD11b+/ Gr-1(−) mouse primary peritoneal (B) and Mouse primary Bone marrow derived Macrophages (C) at indicated time intervals. Cell based CFU analysis clearly indicated that ATLS2021is potent in controlling mycobacterial growth at 9^th^, 24^th^ and 48^th^ hour intervals in these macrophages. Shown here the mean of CFU ± SEM quantified from 3 independent repeats at indicated time intervals

**Figure 5:**
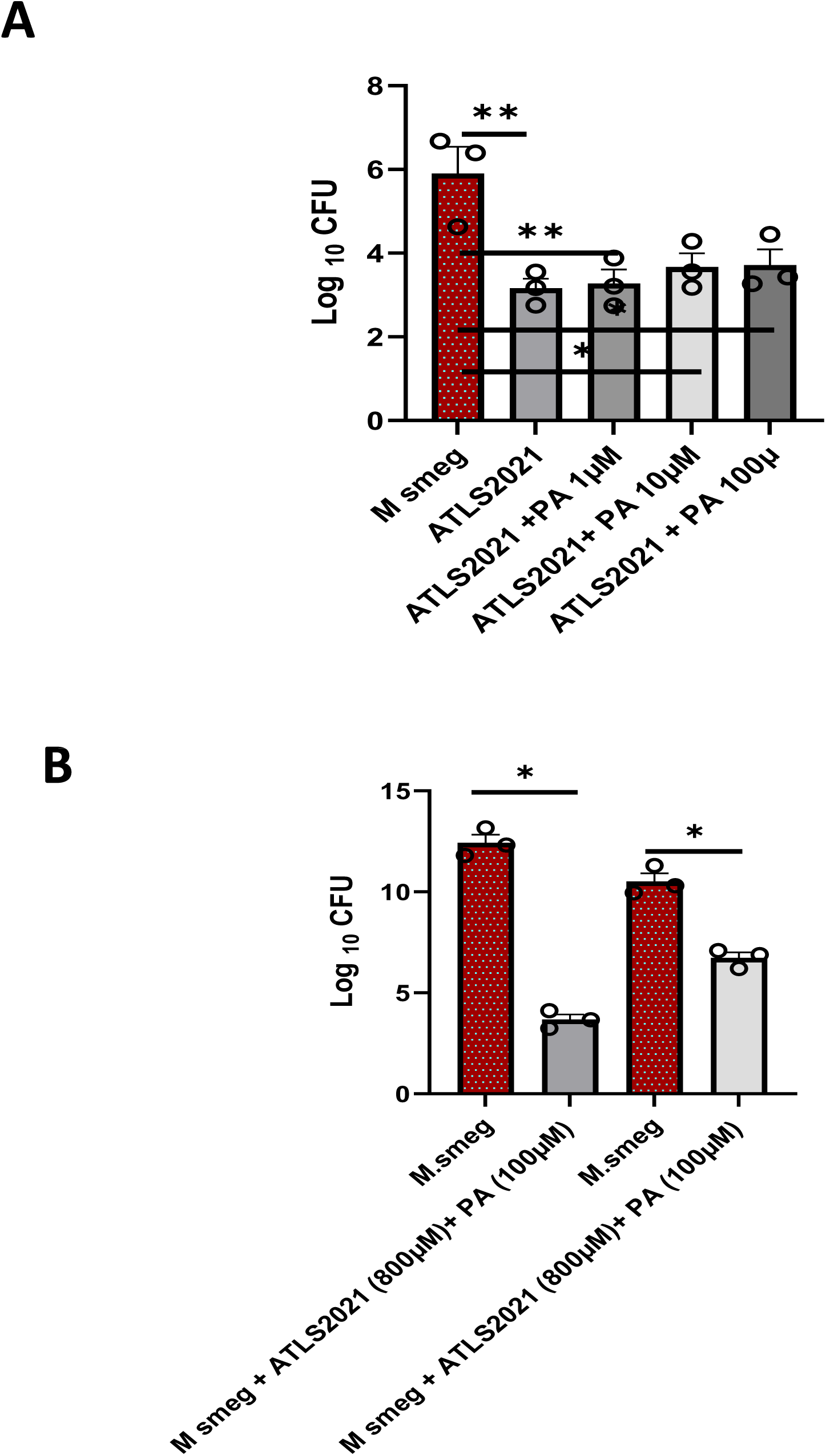
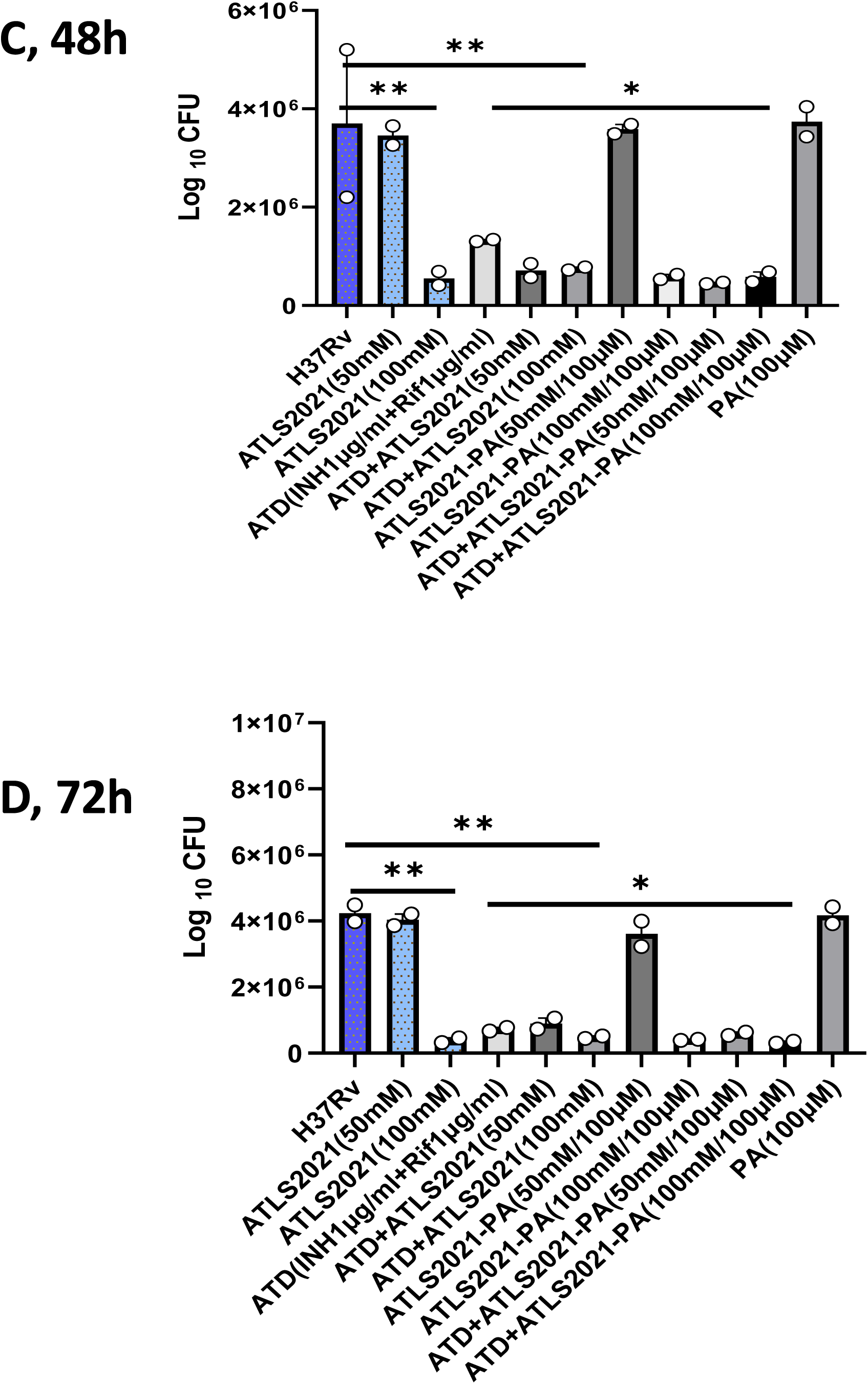
ATLS2021 and PA control mycobacterial growth. Mycobacterial cultures were grown in presence of both ATLS2021+ PA and their growth was accessed as CFU at **24^th^ hour (A)** is broth cultures. The impact of both ATLS2021 and PA was also accessed in RAW macrophages as well. The RAW macrophage were infected with mycobacteria and treated with ATLS2021+PA as indicated and mycobacterial growth was accessed at 48h post infection time interval which clearly indicated anti-tubercular potential of ATLS2021 and PA against *M. smegmatis* **(B)**. In same line, anti-tubercular potential of ATLS2021+PA was also accessed against M tuberculosis (C,D) which in line with *M. smeg*, indicated that ATLS2021 and PA is quite effective in controlling M tuberculosis in RAW macrophages and augment Rifampicin mediated killing of *M. tb* at 48 hours **(C)** and 72 hours **(D).** Shown here the mean of CFU ± SEM quantified from 3 independent repeats at indicated time intervals

**Figure 6:**
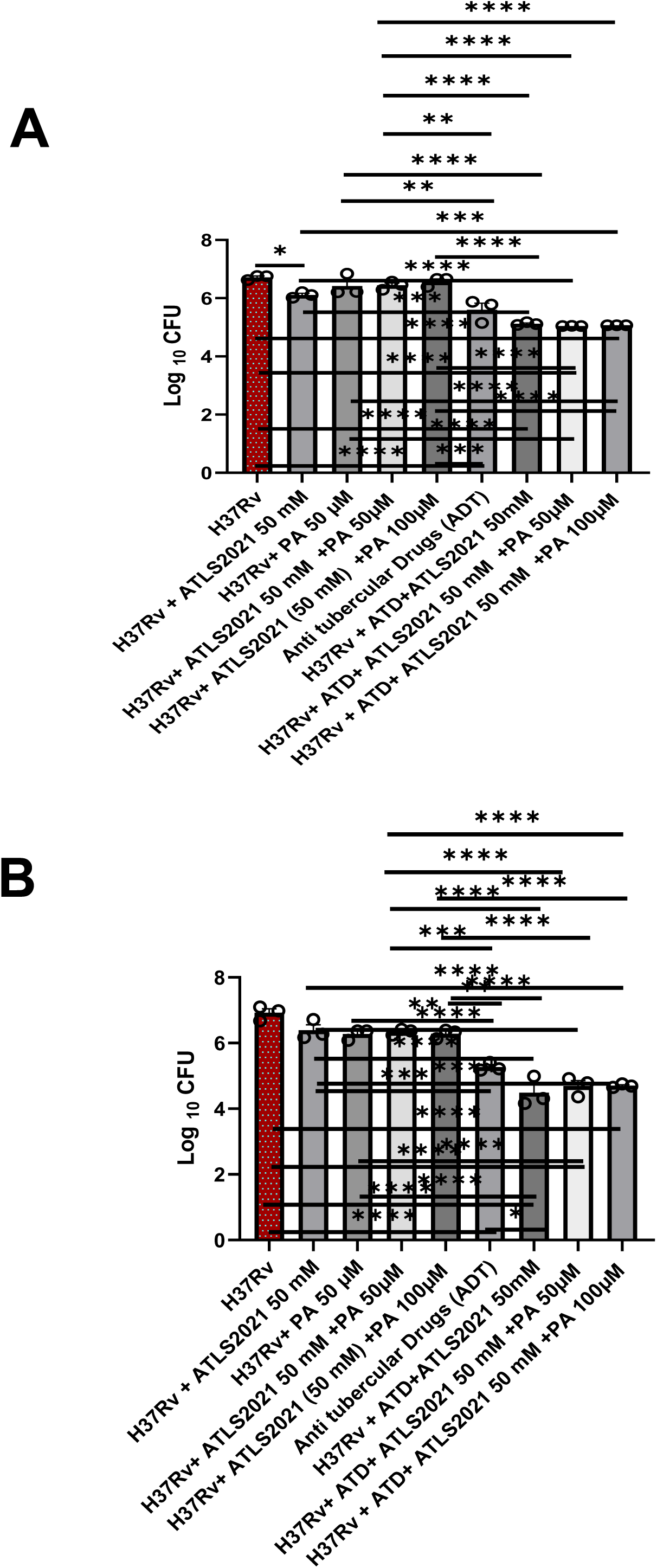
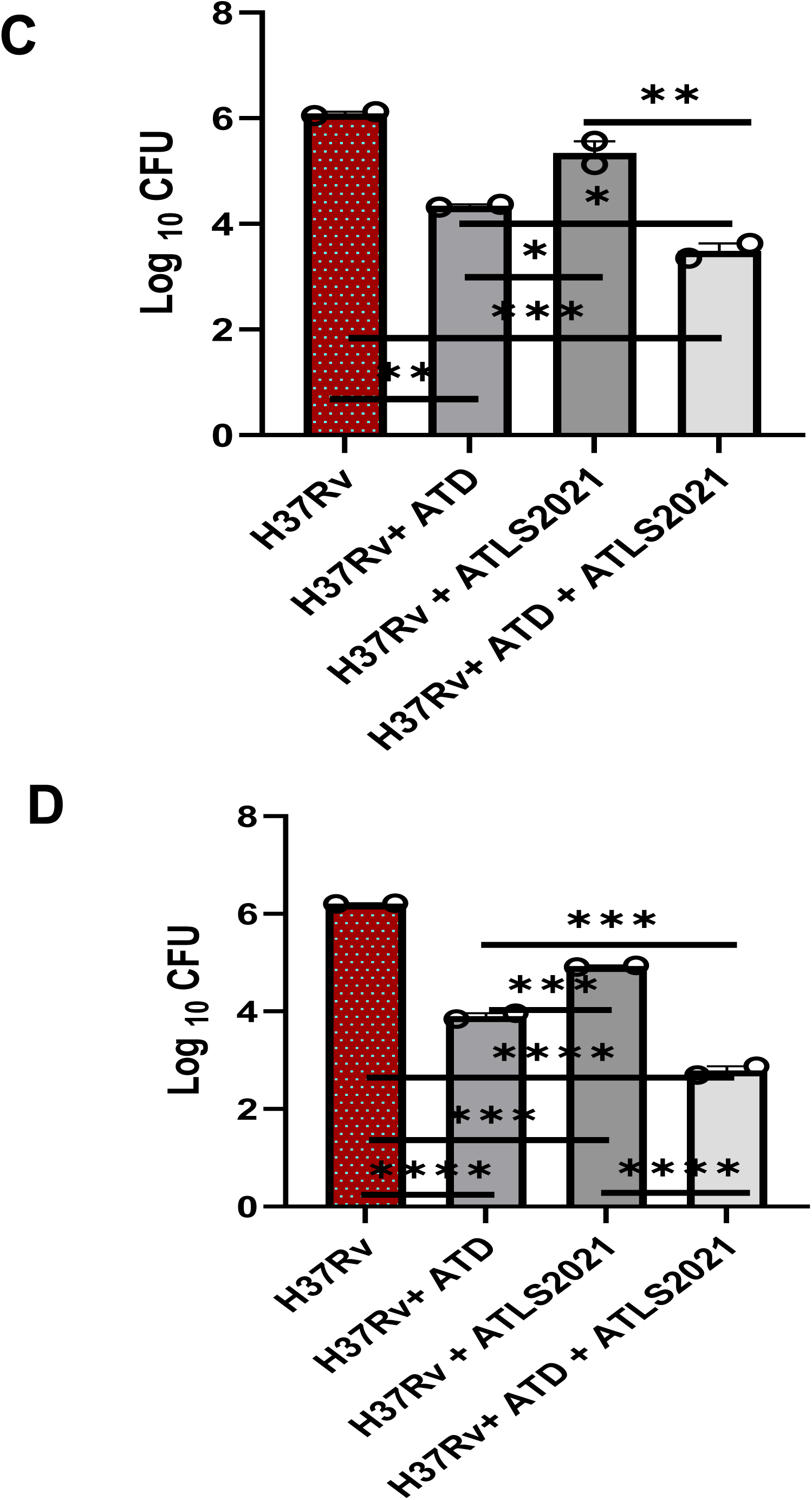
ATLS2021 reduces antimicrobial burden in human monocyte as well. To evaluate the anti-tubercular potential of ATLS2021+PA in human system, anti-tubercular potential of ATLS2021+PA was evaluated in THP-1 human mono/ macrophage cell lines. CFU clearly indicates that ATLS2021 is able to promote killing of *M.tb* in THP-1 mono/macrophages at 48h **(A)** and 72h **(B)** post infection intervals. In same line ATLS2021 was effective on controlling *M.tb* growth in Blood derived CD14+ monocytes at 48h **(C)** and 72h **(D).** Shown here the mean of CFU ± SEM quantified from 2 independent repeats at indicated time intervals.

### ATLS2021PA is able to control *M. tb* burden in the animal model system

Based on anti-mycobacterial impact of ATLS2021 on variety of macrophages line we employed, we were keen to understand whether ATLS2021 would be able to control *M. tb* burden *in vivo* as well. For that purpose, we employed aerosol challenged based study for accessing anti tubercular potential of ATLS2021 in animals. For that purpose, we challenged animals with 1 × 10^7^ (H37Rv) / mouse in aerosol chamber as per scheme shown below **(Figure 7a)** ATLS2021 alone or in combination of PA was administrated both prophylactically and therapeutically as per shown scheme. Anti tubercular drugs (ATD) were used as positive control. After week 8^th^, all mice were sacrificed aspectically and H37Rv growth were monitored in both lungs and spleen by CFU based methods. Following our prediction and in line with *in vitro* results, animal data also demonstrated that ATLS2021 alone or in combination of PA was able to reduce mycobacterial burden significantly in comparism of infection control **(Figure 7b)**. Most interestingly, this remained comparable to ATD group suggesting equi-potent nature of ATLS2021/PA for controlling *M. tb* which was further substantiated by the splenic CFU **(data not shwon).** To further corelate anti tubercular impact with Immunogenic potential of ATLS2021, we quantified pulmonary titre of TNF-α which is most potent anti tubercular in nature. As expected, ATLS2021 supplemented animals had higher titre of TNF-α **(Figure 7c)** in their lungs. Interstingly ATLS2021 enhanced TNF-α titre further in ATD co tratead groups which indicating immunogenic programming potential of ATLS2021 in the infected animals. In same line, we further analyzed the titre of IL-10 which is potentially involved in the clearance of mycobacterial infection. Interestingly supplementation of infected animal with ATLS2021 inhibited the titre of IL-10 **(Figure 7d)** as well as IL-6 **(Figure 7e)** in the lungs of infected animals suggesting the impact of ATLS2021 only on enhanced killing but also improved clearance of infection from the lungs of infected animals.

**Figure 7:**
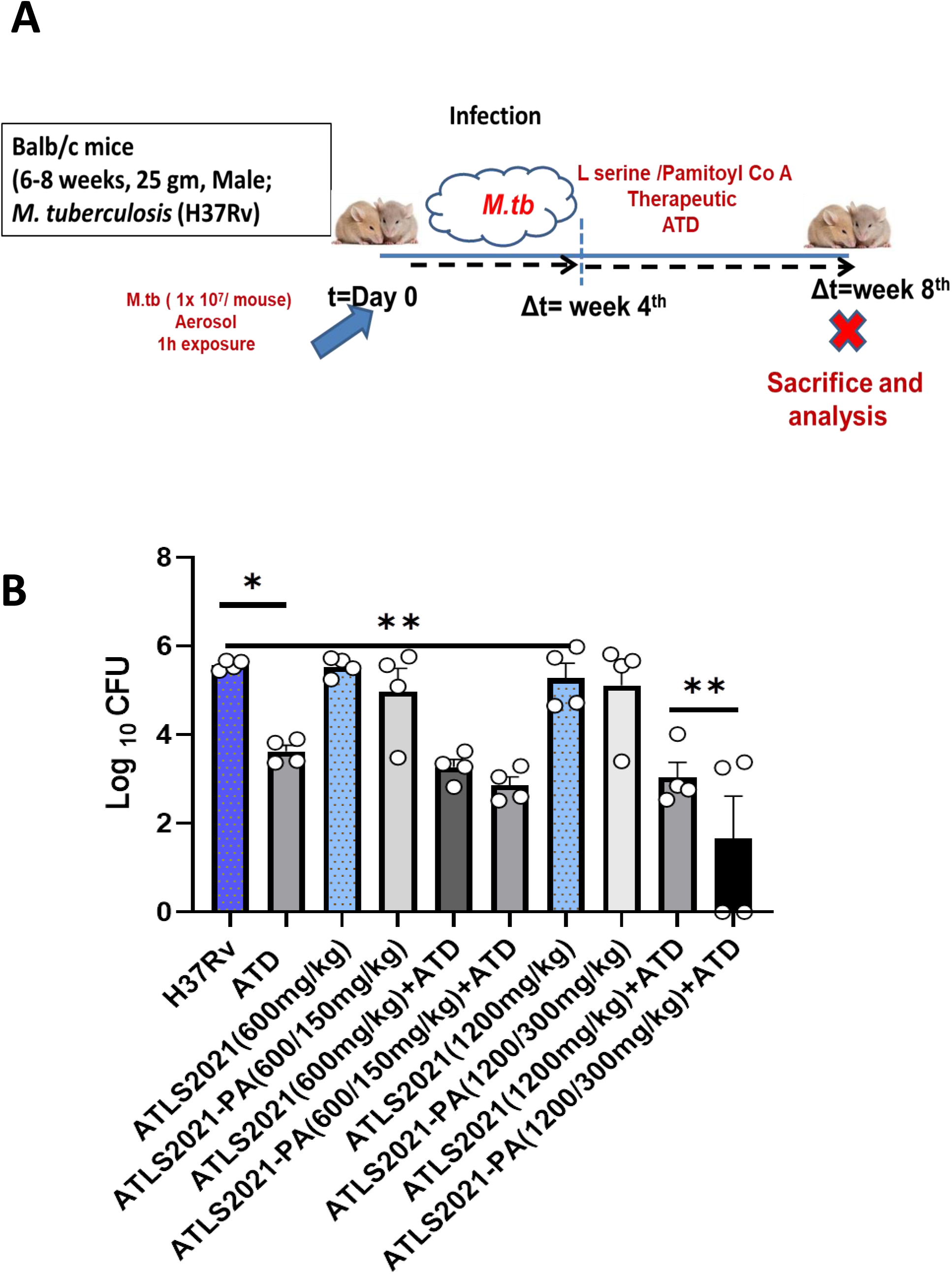
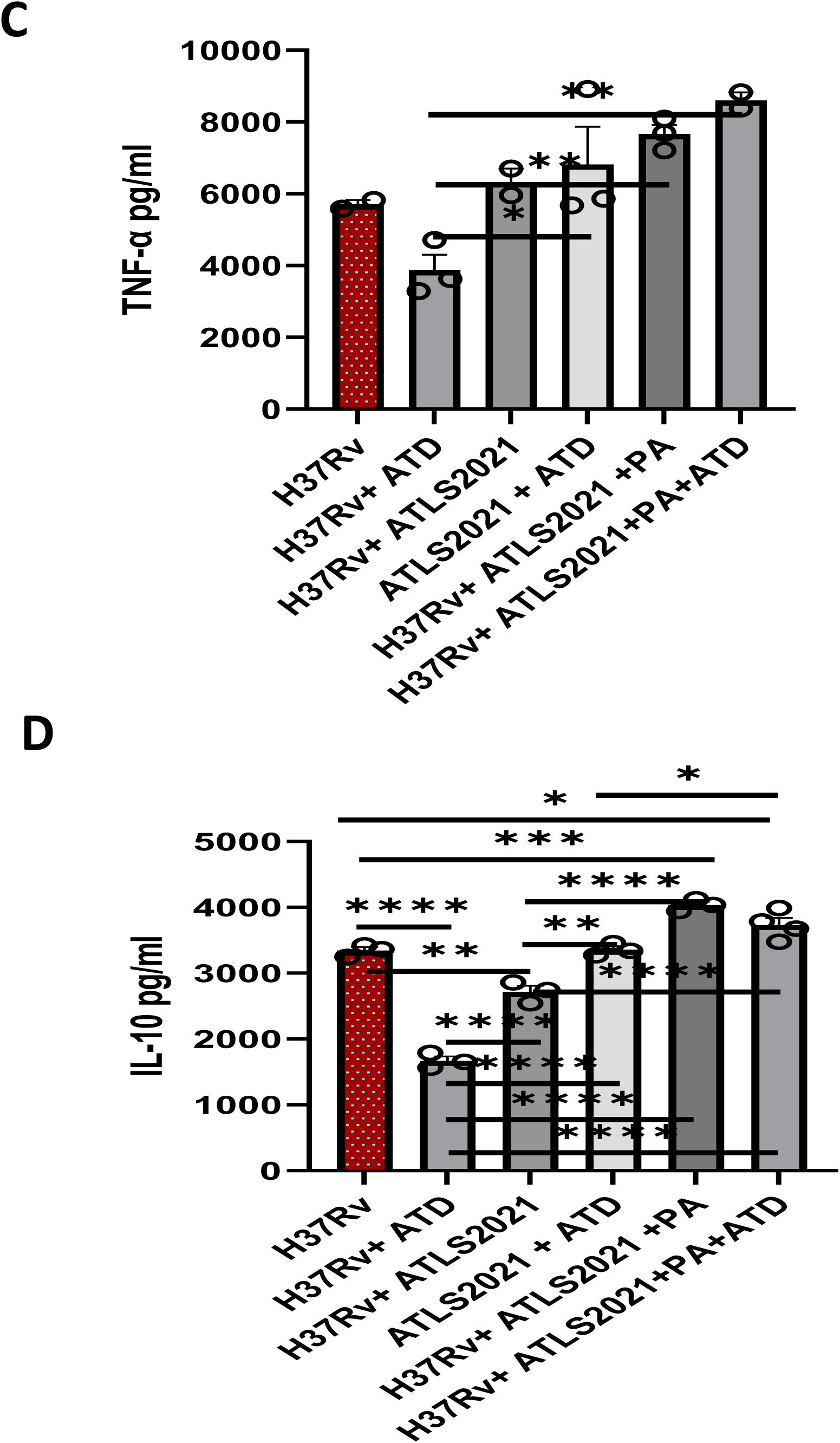
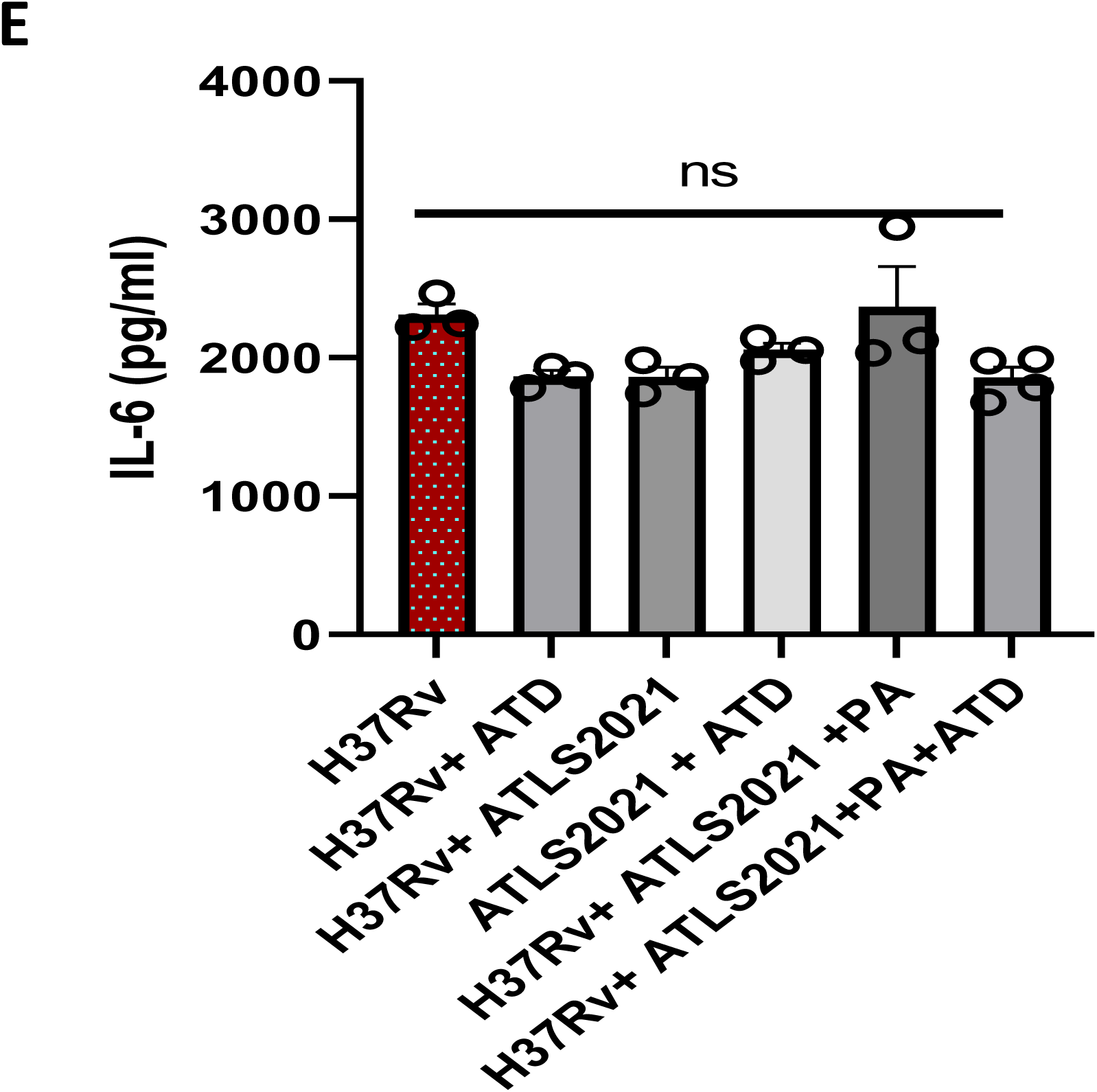
ATLS2021/PA is anti-mycobacterial in the lung of infected animals**. A** Anti-tubercular potential of ATLS2021 was monitored in M tb infected animals as per Scheme shown above. **B**. After completion of 8 week long experiments, animals were sacrificed and their lung homogenate were cultured on LB agar plate and their CFU was calculated after 1 month of incubation. Shown here the mean of CFU ± SEM quantified in various experimental groups. Lung homogenate from various experimental groups were also analysed for the pulmonary titre of TNF-α (C), IL-10 (D) and IL-6 (E). Shown here the mean of *pg/ml* of indicated cytokines ± SEM quantified from various experiments group

### ATLS2021 is potentially immunogenic to murine macrophages

Macrophages utilize both ROS and RNI for killing variety of pathogens including Mycobacteria (12) in host. Our mycobacterial growth pattern analysis clearly indicated anti-mycobacterial role of ATLS2021 which could be attributed to several mechanisms. One of those mechanisms is redox sensing which helps macrophages to respond to various inflammatory signals and respond to the infection for clearing them. On the basis of synergistic impact of ATLS2021 and SNP / CoCl_2_ mediated control of mycobacteria, we predicted that ATLS2021 is probably activating TLR pathways, enhancing NO generation and tweaking phenotypic switch in these macrophages for effective clearance of mycobacteria. In view of the significance of this phenomenon, we assessed the impact of ATLS2021 on NO generation in RAW macrophages. We assessed whether ATLS2021 would enhance the generation of NO in human peripheral blood derived macrophages as well. We anticipated that ATLS2021 may enhance generation of inducible NO in macrophages and to confirm this, we treated RAW Macrophages with various doses of ATLS2021 (which inhibited *M. smegmatis* growth above) and analysed NO titre in the culture supernatant of RAW macrophages. Following our expectation, our results prudently demonstrated that stimulation of RAW macrophages with ATLS2021 enhanced NO in the culture supernatant of naïve **(Figure 8a)** macrophages, Apart from this ATLS2021 augmented ; **LPS (Figure 8b)**; CoCl_2_ **(Suppl Figure 5)** and SNP induced level of NO **(Suppl Figure 5a)** further in the culture supernatant of RAW macrophage This provided evidence that ATLS2021 can augment NO surge in macrophages which are involved in mycobacterial killing by macrophages. We and others have demonstrated that hypoxia promotes mycobacterial killing by RAW macrophages. Hypoxia is known to promote expression of iNOS proteins in macrophages which enhance the generation of NO free radicals and augments bacterial killing in activated macrophages which express iNOS proteins and execute NO dependent killing of intracellular bacteria. To get the hint of this mechanism in our models, system, we treated ATLS2021 supplemented macrophages with CoCl_2_ (HIF-1/ iNOS inducer) and analysed the generation of NO in the culture supernatants of macrophages. Following our prediction, ATLS2021 supplementation enhanced COCl2 induced titter of NO **(Suppl Figure 6)** suggesting the influence of ATLS2021 in activating hypoxic responses in macrophages which is important for clearance of infection and subsequent homeostasis. Apart from clearing pathogen burden, successful antimicrobials should also resolve the inflammatory response. In line with this hypothesis, ATLS2021 inhibited *M. smegmatis* induced secretion of TNF-α and IFN-γ cytokines in RAW **(Suppl Figure 7), CD11b+ / Gr-1**(**−**) mouse peritoneal **(Suppl Figure 8), CD11b+ / Gr-1**(**−**) bone marrow derived macrophages **(Suppl Figure 9)** revealed homeostasis potential of ATLS2021 while controlling mycobacterial burden. Anti-tubercular and immunogenic potential of ATLS2021 for MDR TB in both murine and human macrophages provoked us further to test whether ATLS2021 would condition blood derived CD14+ monocytes from MDR PTB patients as well. To demonstrate this CD14+ monocytes were purified as per method described earlier and supplemented with ATLS2021 and titre of NO and indicated cytokines were quantified. In line with murine macrophages, ATLS2021 enhanced generation of NO constitutively **(Figure 9a)** and augmented LPS **(Figure 9b)** induced NO levels. Most surprisingly ATS20221 augmented even rifampicin induced NO level also **(Figure 9c)** in CD14+ monocytes from healthy donors as well to our surprise from MDR TB patients suggesting its immunogenic potential on CD14+ macrophages. This observation was further substantiated by augmentation of ATLS2021 mediated IFN gamma titre **(Figure 9d) (Figure 6b)**. Concomitantly, ATLS2021 inhibited the secretion of both IL-10 **(Figure 9e)** & IL-6 titre **(Figure 9f)** particularly in CD14+ macrophages from MDR TB patients indicating immune conditioning potential of ATLS2021 CD14+ monocytes MDR TB patients. .Quite convincingly ATLS2021 enhanced the gene signature of all key enzymes which are involved in the biosynthesis of Oxysphingolipids in CD14+ macrophages from both healthy as well as MDR patients. Infect inhibition of Sphk-2 and SMAseD mRNA **(Suppl Figure 10)** indicting a bias of ATLS2021 for enhancing Sphingomyelin-ceramide pathways which are involved in immunogenic inflammation and anti-mycobacterial response in host (3). To further correlate this with anti-mycobacterial potential of ATLS2021, we tested whether ATLS2021 would be effective in controlling MDR TB in macrophages or not. Indeed, following our prediction, our preliminary growth pattern analysis of (SNMC/90) revealed that ATLS2021 is capable of controlling burden of MDR TB as well in RAW macrophages and nicely supported immnne-conditioning potential of ATLS2021 on macrophage. These results altogether highlighting its potential against MDR TB as well **(Figure 9g & 9h)** inducting its translational potential as effective anti tubercular rgime having immune condtionning potential for managing MDR TB.

**Figure 8:**
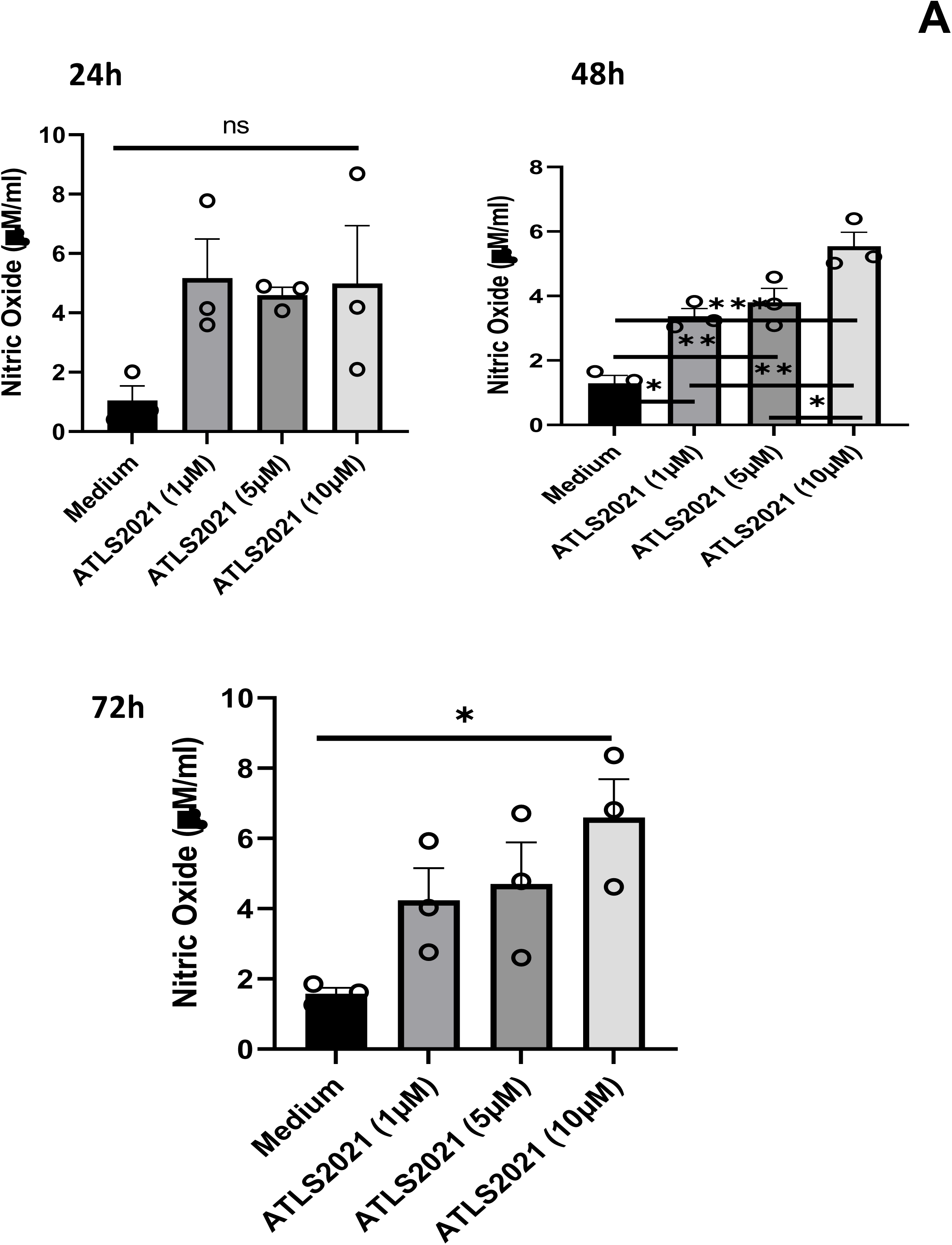
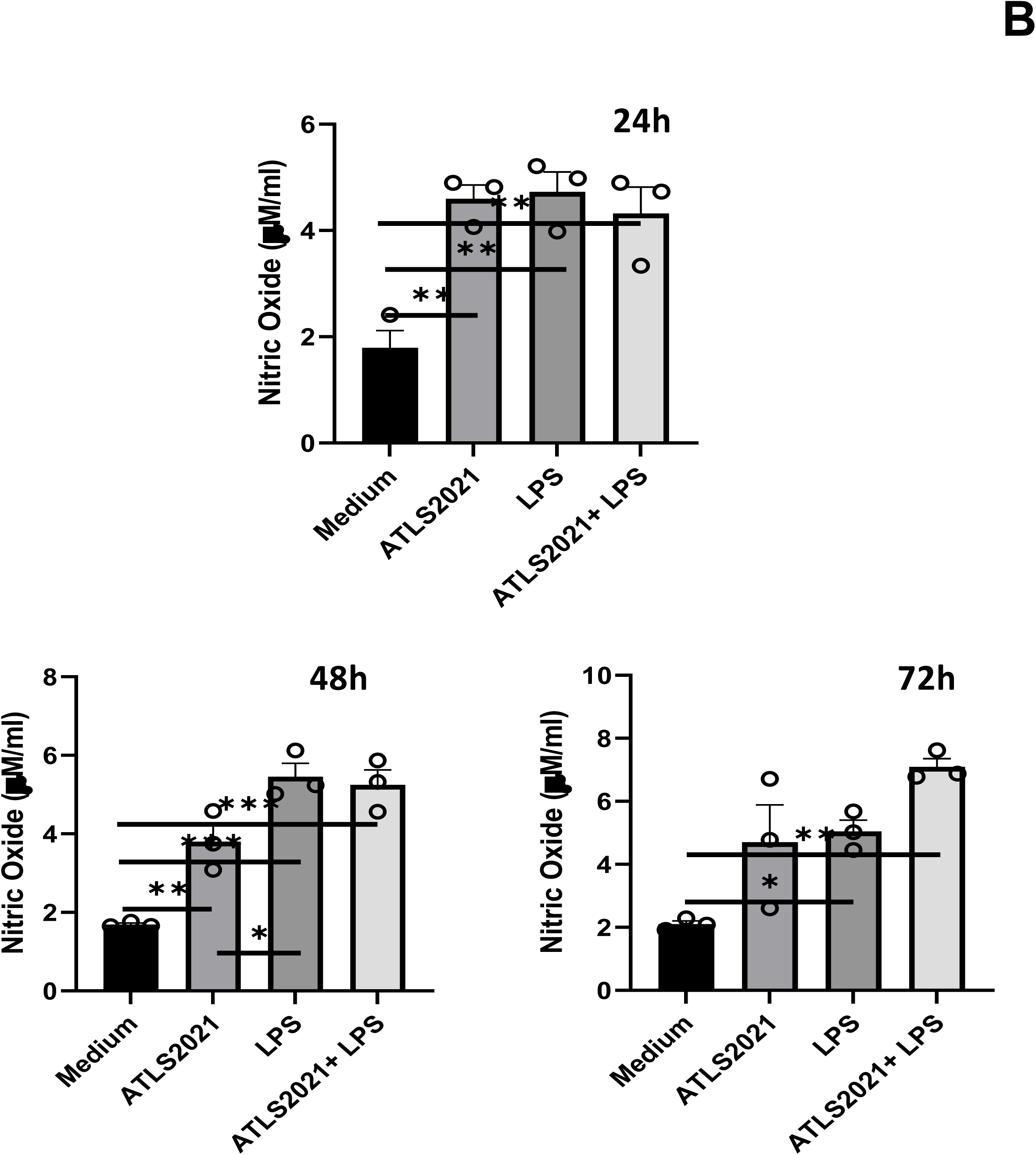
ATLS2021 potentially activate macrophages. RAW macrophages were treated with ATLS2021+PA alone **(A)** or in combination of LPS **(B) and** Nitric oxide levels were quantified in the culture supernatant of RAW 264.7A macrophages. Shown here the mean µM of NO ± SEM quantified from 4 independent experiments.

**Figure 9:**
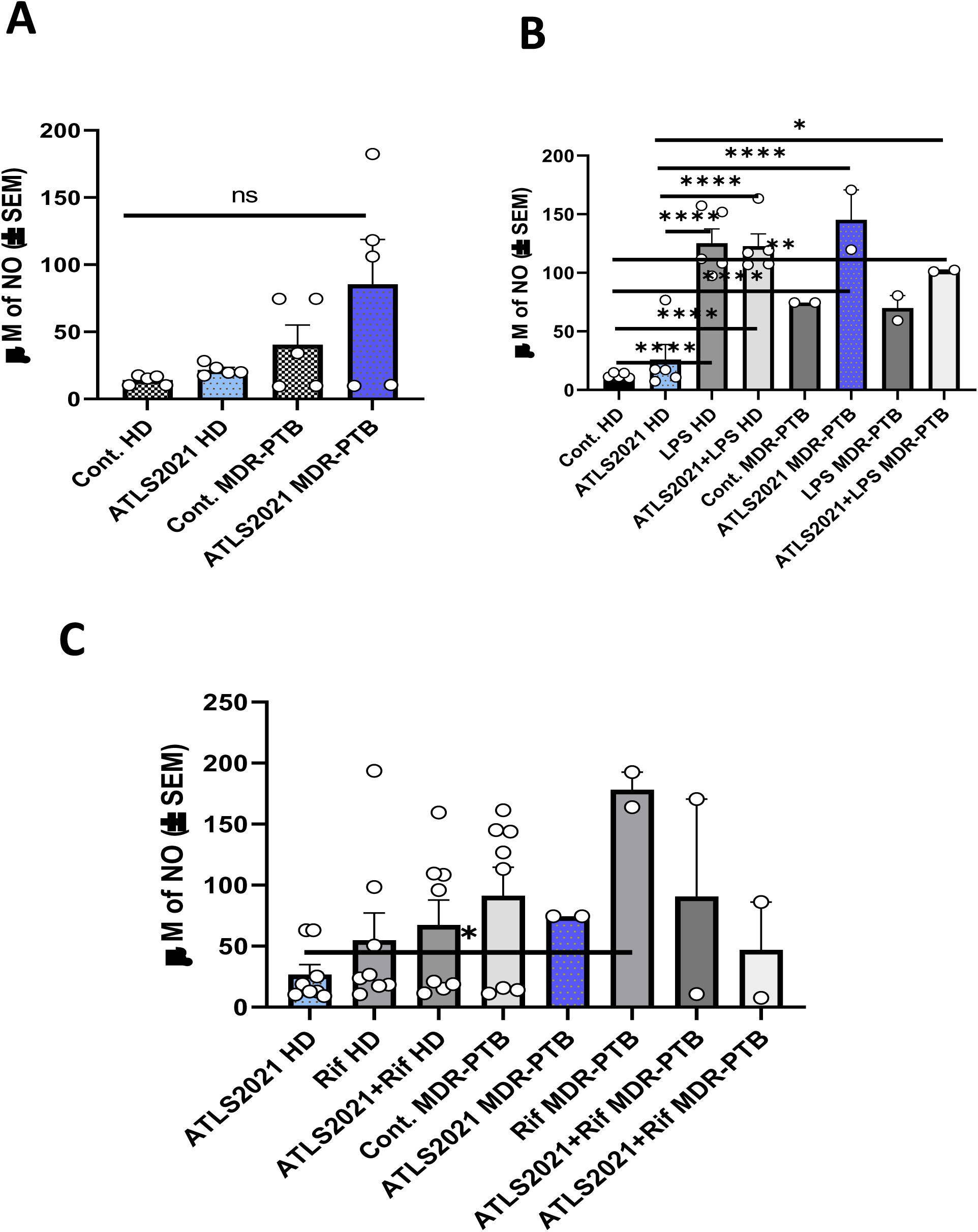
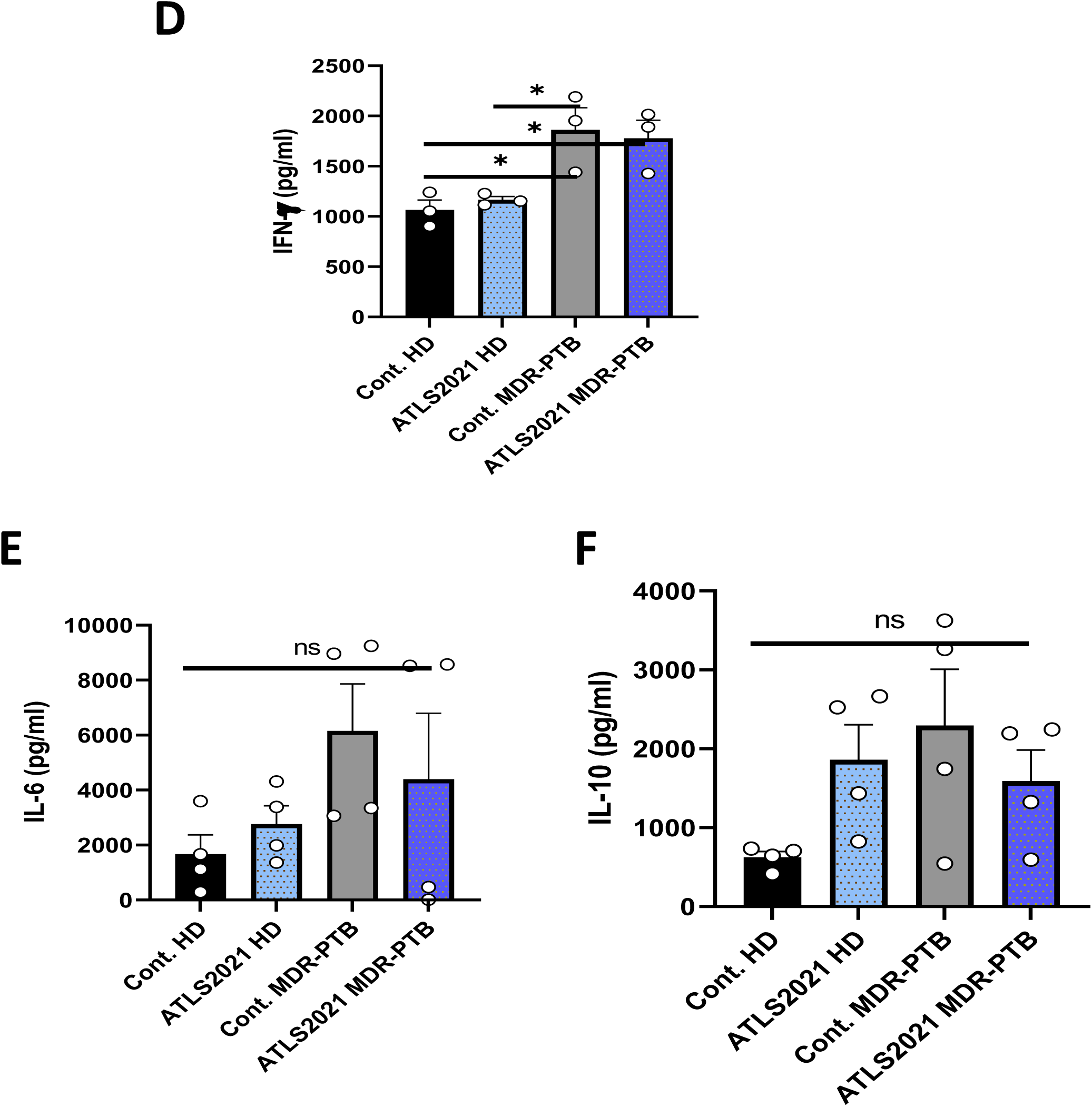
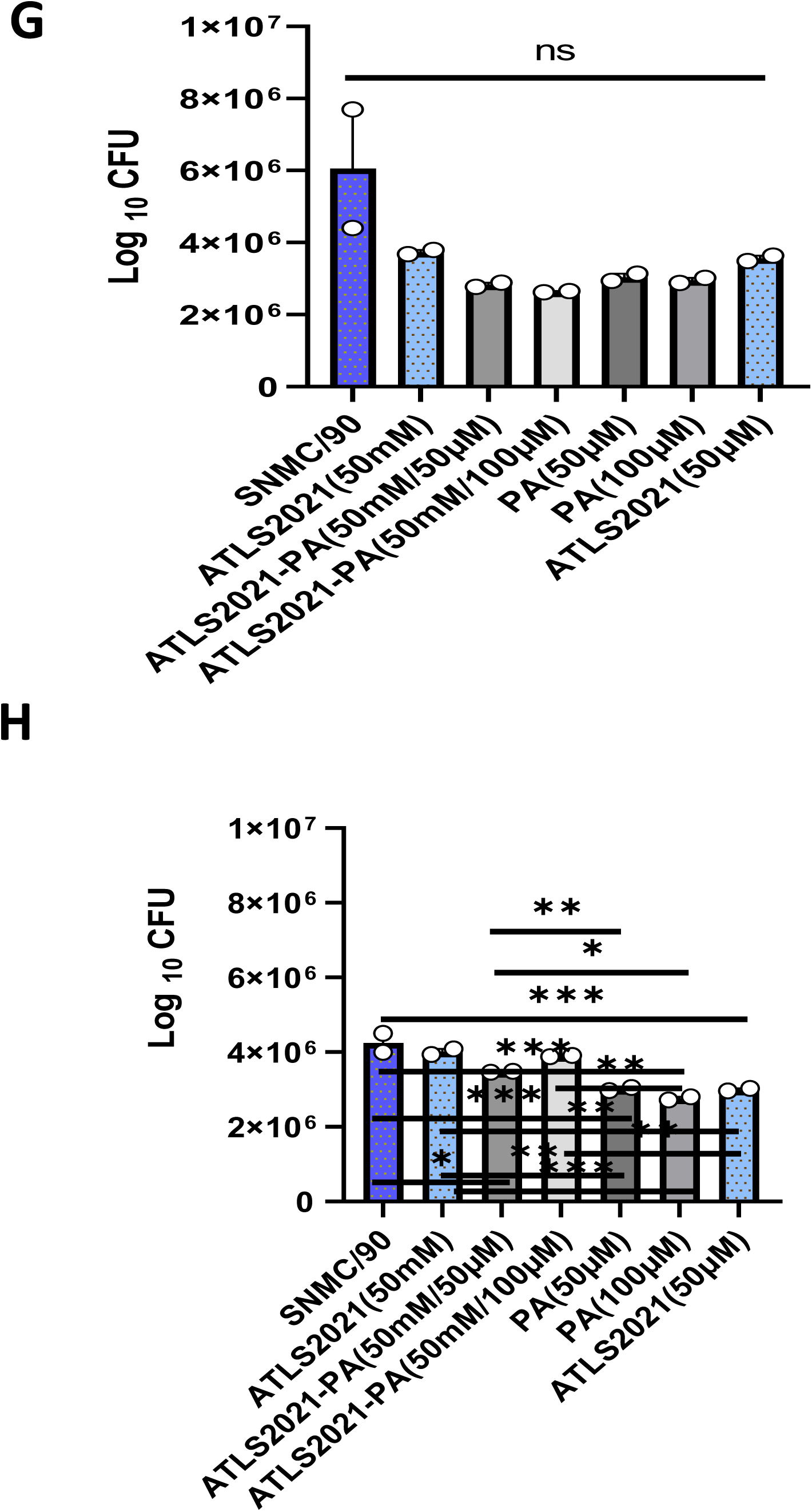
ATLS2021 tweak innate immune response in CD14+ human macrophages. CD14+ monocyte were purified from blood form healthy and PTB MDR TB patients. These cells were 95% pure at the time of cultures. These cells were treated with ATLS 2021 along with LPS and Rifampicin and NO titres were quantified at 48h post treatment **(A-C).** Shown here the mean µM of NO ± SEM quantified from CD14+ monocytes obtained from 5 healthy and MDR PTB patients. The culture supernatant of these cells were also analysed for the secretion of IFN-ɣ **(D)**, IL-6 **(E)** and IL-10 **(F)** at 48h post stimulation. Shown here the mean of *pg/ml* of indicated cytokines ± SEM quantified from CD14+ monocytes obtained from 5 healthy and MDR PTB patients. G-H. To correlate ATLS2021 induced innate immunity in CD14+ monocytes from MDR TB patients, we accessed anti tubercular potential of ATLS2021+PA in RAW macrophages. These cells were infected with clinical isolate of MDR TB and CFU were quantified at 48 h **(G)** and 72 h **(H)**. Shown here the mean of CFU ± SEM quantified from 2 independent repeats at indicated time intervals

## Discussion

Metabolic programming of host limits effective control of variety of pathogens by host (13). Most of TB patients’ cohort is poor in their nutrition / metabolic index (low Sphingolipid content in their serum in particular) (14) and suffer from severe disease. We here report that immune metabolic programing of host (15) (16) has decisive impact on the management of TB. Several strategies including antibiotics/ dietary supplementation and other immunogenic approach are being tailored for augmenting immunogenic inflammation for controlling the mycobacterial burden in host. However several evidences have suggested that Mycobacteria subvert host immune response in securing survival in hostile environment of host. MDR TB in particular, is the typical example which is trained to thrive in pathogenic inflammation micromilieu of lung (17) which they establish in host for their opportunistic survival (18). Host and mycobacteria compete for their nutritional demand and secure their respective survival. In most of drug resistant cases which are not able to control the infection, immune metabolic programing of macrophages in particular is one of the pathophysiological manifestations (19) which predispose host sensitive for pathogen burden. Several nutrients, certain amino acid and lipids are involved in tweaking immunogenic response in host (20) for protective immunity of host against mycobacteria. In this context our study potentially advocates us that supplementing host with homeostatic nutrients likes ATLS2021PA is certainly helpful to the host for controlling mycobacterial burden. On the basis of its remarkable normalization of neuropathic diseases (Ref clinical trials) and our own data, it is evident that tweaking host Sphingolipid pathways either via supplements and bio-therapeutics means (ATLS2021PA) would condition infected lungs immunologically for effective clearance of mycobacterial infection. This approach has been patented by us in past and currently under clinical trial and commercialization phase.

The approach which we have adopted in our study holds tremendous therapeutic potential and believed to contribute significantly to existing host directed therapy against TB. This proposal would be delivering new category of treatment modality for controlling *M. tb* burden much more effectively. We believe that such immune-mediated interventions would be helpful in managing patients with latent tuberculosis by fine tuning their metabolic demands. We also believe that inclusion of ATLS2021PA with currently used ATD would certainly change the treatment modality for both active (potentially) and latent patients (quite possibly) for effective clearance of the infection burden in patients. The major scope of this study is to establish enhanced level of Oxy-Sphingolipids which was evident by our real time PCR data and encounter mycobacterial persistency related aberrant immuno-pathologies like ARDS or Pulmonary Fibrosis. This proposal would be delivering new adjunct therapy for controlling *M. tb* burden much more effectively. We believe that such immune-mediated interventions would be helpful in managing patients with latent TB which need additional approaches.

## Material and Methods

### Antibodies and Reagents

All reagents were purchased from Sigma-Aldrich (UK), unless stated otherwise. RPMI 1640, Penicillin Streptomycin solutions were procured from the Sigma-Aldrich (Cat. No?). Recombinant mouse IFN-γ cytokine was purchased from eBiosciences, (San Diego, CA). CD11b+ human and Mouse MACS Microbeads and LC Columns were purchased from Miltenyi Biotec limited. ATLS2021PA was purchased from Sigma. TNF-α, IFN-γ and IL-6 ELISA kits were purchased from R&D system (Darmstadt, Germany). Human AB serum was purchased from (H4522-Sigma-Aldrich) 7H9 and 7H10 was purchased from Difco (Cat. No?). Real time PCR primers for Sphingolipids pathways were synthesized from (Eurofins, Genomics). RNA easy kit was purchased from (Qiagen, QIAwave RNA Mini Kit) SYBR Green was purchased from (BIORAD, SYBR^®^ Green Master Mix cat no.?) Rifampicin (SBR00067-Merck) and Isoniazid (I3377-Merck) ani tubercular drugs were purchased from Merck. FBS was purchased from (12103C-Sigma-Aldrich) Ficoll hypaque was purchased from (Histopaque, 10771, Sigma-Aldrich).*M. smegmatis*, *H37Rv* cultures were purchased from MTCC IMTECH Chandigarh while clinical isolate of MTB was isolated from MDR patients from SN Medical Hospital, Agra. SNP and CoCl_2_ was purchased from Merck(Cat. No.?). All plastic wares were purchased from (Genaxy, Trueline). Sulphanilamide and N-(naphthyl) ethylene-diamine**-** dihydrochloride were purchased from Sigma. REMA analysis was done as per manufacturer protocol. Resazurin dye was purchased from (R7017**-** Sigma-Aldrich).

### Ethical approval

The entire study protocol was approved by the Institute Biosafety, Animal and Human Ethics Committee of Amity University, Noida, JALMA institute Agra and AIIMS New Delhi. Collection of blood from healthy donors at Amity University was approved by IEC of Amity University (approval number viz AUUP/IEC/May 2023/7 IEC). Animal experiments at amity were approved by Institutional Animal Ethical committee (approval number-AUUP/AIP/16/22/01-04). The animal experiments at JALMA were approved by their IAEC wide their approval number – NJIL&OMD /61AEC/2022-05. Prospective clinical study with TB patients was approved by IEC of AIIMS wide their approval number AIIMS/REV2023-44. Written Informed consent was obtained from healthy donors and PTB patients before blood collection.

### REMA analysis

To estimate the MIC of Rifampicin for we performed REMA analysis. For that purpose, two-fold serial dilution of ATLS2021 ranging from 0.625–2.5 mM, and 1.5-50 µg/ml of rifampicin were added in the wells. The *M. smegmatis* MTCC 9944 bacterial suspension of No.1 McFarland standard (∼3.2×10^6^cfu/ml) was prepared and diluted 1:100 in 7H9 broth. 100µl inoculums were aliquoted into each well of a 96-well plate. Perimeter wells of the plate were filled with sterile water to avoid dehydration of medium during incubation as per table no-1 −3. After two days of incubation, Resazurin dye was added to each well, sealed and re-incubated at 37 °C for overnight. The visual MIC was defined as the lowest drug concentration that prevented the colour change for Resazurin reagent from blue to pink. Blue colour in the well was interpreted as there is no mycobacterial growth and pink colour was scored as growth occurrence. Additionally, treated or untreated cultures from 96-well plates were serial diluted and plated on 7H11 agar plate to determine the bacterial viability. The addition of 5 mM ATLS2021 had no significant effect on the survival of mycobacteria. The MIC of rifampicin against *M. smegmatis* was 25µg/ml. Addition of ATLS2021 at 0.625-2.5mM in the presence of rifampicin did not completely inhibit the growth of mycobacteria. 100 µl of the treated culture from a parallel experiment was serially diluted and plated on M7H11 agar. The result showed that 2.5mM potentiated the rifampicin effect. Treatment with ½ MIC of rifampicin (12.5 µg/ml) decreased the bacterial survival by 10 folds i.e. from 5X 10^10^ to 3 X 10^9^ CFU/ml. Addition of ATLS2021 at 2.5mM potentiated the activity of rifampicin and the bacterial survival decreased further by 100-fold i.e., from 5X 10^10^ to 4 X 10^8^ CFU/ml To evaluate this, 50 µl each of ATLS2021 and rifampicin were added into 100µl of *M. tb* bacterial culture (1:20) in each well of a 96-well plate using the Resazurin dye method as described above by the checkerboard assay. The plates were then incubated at 37 °C after for 7 days. The Resazurin dye was added, and plate was further incubated for 1 more day. In the Checkerboard technique, the interaction between ATLS2021 and rifampicin against *M. tb* were predominantly additive (ΣFIC for ATLS2021 with rifampicin=0.78 mM) indicating the activity of ATLS2021 with rifampicin combination being greater to the sum of their independent activity.

### Cell Culture

RAW 264.7 macrophages and THP-1 monocytic cell line were purchased from NCCS Pune and maintained in Roswell Park Memorial Institute medium (RPMI) containing amino acids and supplemented with 10% (v/v) fetal bovine serum (FBS) and 1% penicillin and streptomycin in CO_2_ incubator. To isolate CD11b+ peritoneal macrophages, C57BL/6j mice were injected with 1ml of 4% brewer thioglycolate medium intraperitoneally and peritoneal lavage was harvested at third day post injection. Peritoneal lavage was centrifuged at 400g for 8min and the cell pellet was re-suspended in fresh serum free RPMI. CD11b+ macrophages were purified by MACS-based separation method and were cultured in serum containing medium overnight. Macrophage monolayers were washed on the following day. For the preparation of Bone Marrow-derived Macrophage (BMDM), C57BL/6j mice were sacrificed by cervical dislocation and both femurs were excised aseptically. Femurs were washed twice with ice-cold sterile PBS. Tibia were cut from the femur at the joint and purged with ice cold PBS using 5 ml syringes. The cell suspension were collected in 15 ml tubes and filtered through a 70 μm cell strainer to remove cellular debris. The cell suspensions were centrifuged at 400g for 10 min at 4 °C. RBC in pellet were lysed by using ACK lysis buffer and removed by centrifugation. Cell pellets were dissolved in complete DMEM and incubated in presence of GM-CSF for a week time to obtain BMDM, supernatants were discarded and the attached macrophages were washed and detached by gentle pipetting the PBS across the dish. Cells collected were centrifuged at 1200 rpm for 5 minutes and re-suspended in 10 ml of BMDM cultivation media containing 10% fetal bovine serum and 2 mM L-glutamine. Cells were counted, seeded and cultivated in tissue culture plates 12 hours for attachment before the experimentation.

### Purification of blood derived CD14+ monocyte from healthy donors and PTB patients

A total of 10 new PTB patients with age groups from 18-55 years that were displaying clear clinical symptoms, such as benchmark respiratory problems, positive mucus smear microscopy, culture tests, and radiology of chest, were recruited from the Outpatient Department (OPD) of AIIMS, New Delhi. For purifying CD14+ monocytes from blood of healthy and PTB patients, written consent was obtained from participants before sample collection. A total of 10 new PTB patients with age groups from 18-55 years that were displaying clear clinical symptoms, such as benchmark respiratory problems, positive mucus smear microscopy, culture tests, and radiology of chest, were recruited from the Outpatient Department (OPD) of AIIMS, New Delhi. A total of 10 healthy individuals with age group from 18-55 were recruited. Healthy group individuals with no history of TB, mycobacterial infections, or other infectious diseases and PPD negative were included in the study. For purification of CD14+ monocyte from blood, 10 ml of blood sample were layered over ficoll and tubes were centrifuged at 2000 rpm for 20 minute at 25°C with break free. Subsequently the PBMC were collected from the ring and washed with PBS 3X at 1500 rpm for 10 minute each. These PBMC were counted and incubated with CD14+ magnetic beads for 10 min on ice. The labelled PMBC were passed through LD column under high magnetic field and CD14+ monocyte were purified using positive selection method. The cells were washed and counted and their purity was checked by FACS which was found 95%. These cells were cultured over night for rest and stimulated with ATLS2021 for 48h intervals and their culture supernatant were collected for NO and cytokines analysis.

### Cellular infection

RAW264.7, peritoneal and Bone marrow derived macrophage and CD14+ monocyte from healthy donors were seeded in 24-well tissue culture plates (0.2×10^6^/well) and incubated overnight in CO2 incubator at 37 °C. On the following day, cultured macrophages were infected with cultures of H37Rv. Bacterial clumps were remove by passing culture through needle 15-20 times. After 3h of infection, macrophages were washed thrice with PBS and then cultured in Gentamycin (10µg/ml) containing medium to remove extracellular bacteria. Cells were incubated further and bacterial growth was analyzed by plate based method. Cell culture supernatants were collected for NO quantification. To demonstrate bacterial killing by macrophages, bacterial counts were performed over a time period of 1, 4 and 7 days. For this purpose cell lysates were serially diluted in sterile PBS and different dilutions were plated over Middle-brook 7H10 agar plates supplemented with OADC and Tween80. The bacterial colonies were counted documented as colony forming units (CFU) per milliliter.

### *M. tuberculosis* infection of mice

For animal infection, frozen aliquots of *M.tb* (H37Rv) were thawed, washed in phosphate buffer saline (PBS). For *in vivo* studies, C57BL6mice were purchased from NIN, Hyderabad, were maintained at 20–22°C and relative humidity of 50–70%. *M. tb* infection experiments were conducted at BSL-3 facility of JALMA, Agra, India. Six to eight week old mice were kept in well-ventilated perplex boxes and the mice were given standard diet of rodent pellets and water. Each mouse was challenged with M. tuberculosis (H37Rv) via respiratory route using an aerosol chamber (Inhalation exposure system, Glasscol Inc., IN, USA) for 45 min in 100 µl of saline. Mice were treated with ATLS2021 and PA two weeks after infection in therapeutic setting weekly for the duration of incubation. After infection, mice were kept in controlled environment and their health status was monitored regularly. Lungs were weighed, placed in 1 ml of PBS solution in sterile tubes, chopped with sterile scissors into small pieces and homogenized. Tissue homogenates were centrifuged at 4,000g and the supernatants were filtered through 0.22μm filter. Cytokines were quantified using sandwich ELISA kit and concentration of each cytokine was quantified using standard curve according to manufacturer’s instructions.

### Macrophages stimulation analysis

RAW264.7 macrophage cell lines, mouse peritoneal and bone marrow derived macrophages and blood derived CD14+ monocytes were infected with mycobacteria in presence of ATLS2021 and various other stimuli used and their innate immune response were accessed by quantifying NO and Cytokines titres by Griess reagent and ELISA method respectively. Level of Nitric oxide levels in macrophage culture supernatants in various experimental groups was quantified as nitrate by standard Griess reagent method. Equal volumes of the culture supernatants and Griess reagent (1% sulphanilamide/0.1% N-(naphthyl) ethylene-diamine-dihydrochloride prepared in 5% o-phosphoric acid) were mixed and incubated. Absorbance was recorded at 550 nm by spectrophotometer. NO titres in samples were quantified against a NaNO_2_ standard curve generated For quantification of cytokine secretion from above from above cultures were collected after 24hrs, 48hrs and titre of Il-10, IFN, TNF and IL-6 cytokines were estimated by using commercially available kits from R & D Systems, USA.

### Statistical Analysis

Data was statistically analysed by using GraphPad Prism 3 and STATA, USA software. Difference between the groups was calculated by Mann Whitney test and p value less than 0.05 was considered as significant. One-way ANOVA test was used when more than 2 groups were compared. Student’s unpaired, 2-tailed t test was used when comparing 2 groups.

## Acknowledgment

This work was supported by a grant from the ICMR grants to HP and AKS. We thank all the study participants and the staff of the Staff of ICMR-NJIL&OMD for helping us in the animal challenge experiments

## Conflict of Interest

This approach is already patented and under commercialization by us.

## Supplemental data

**Supplementary Figure 1:**
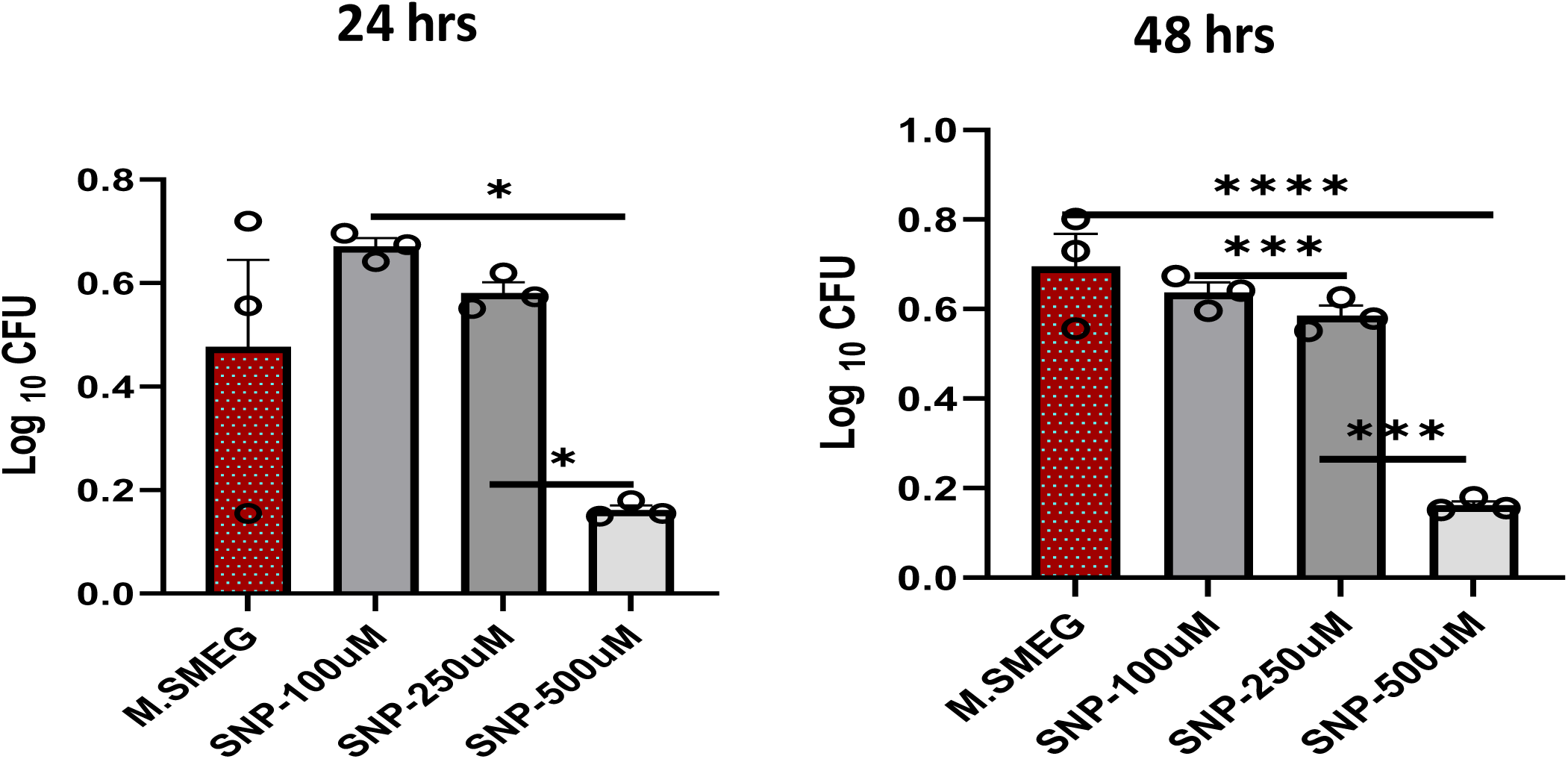
SNP reduces antimicrobial burden. Mycobacteria were cultured in presence of varying concentration of SNP and their CFU was calculated at 24 hours and 48 hours. Shown here the mean of CFU ± SEM quantified from 2 independent repeats at indicated time intervals

**Supplementary Figure 2:**
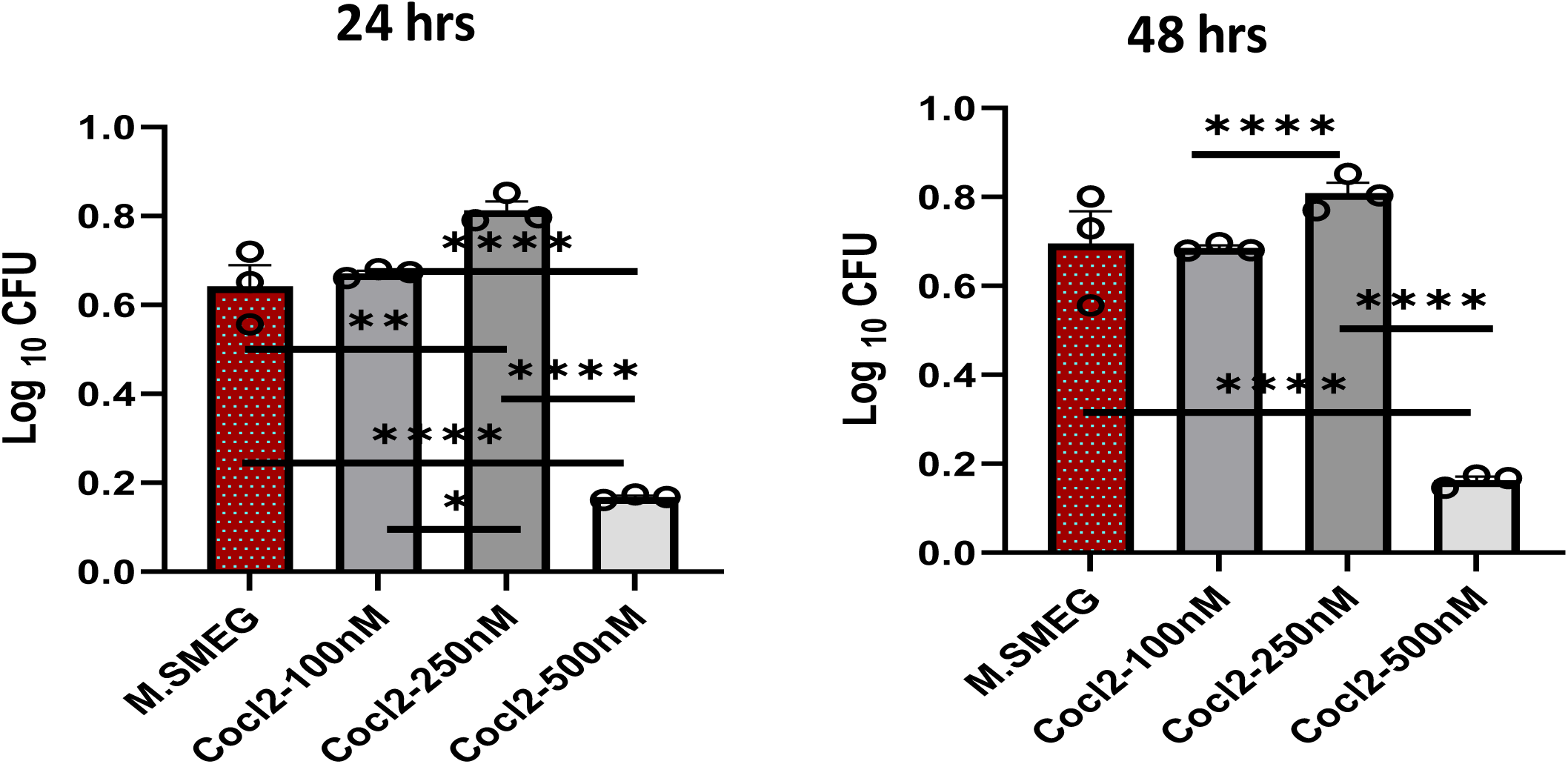
Hypoxia mimetic Cocl2 potentially reduces antimicrobial burden. Mycobacteria were cultured in presence of varying concentration of CoCl2 and their CFU was calculated at 24 hours and 48 hours. Shown here the mean of CFU ± SEM quantified from 3 independent repeats at indicated time intervals

**Supplementary Figure 3:**
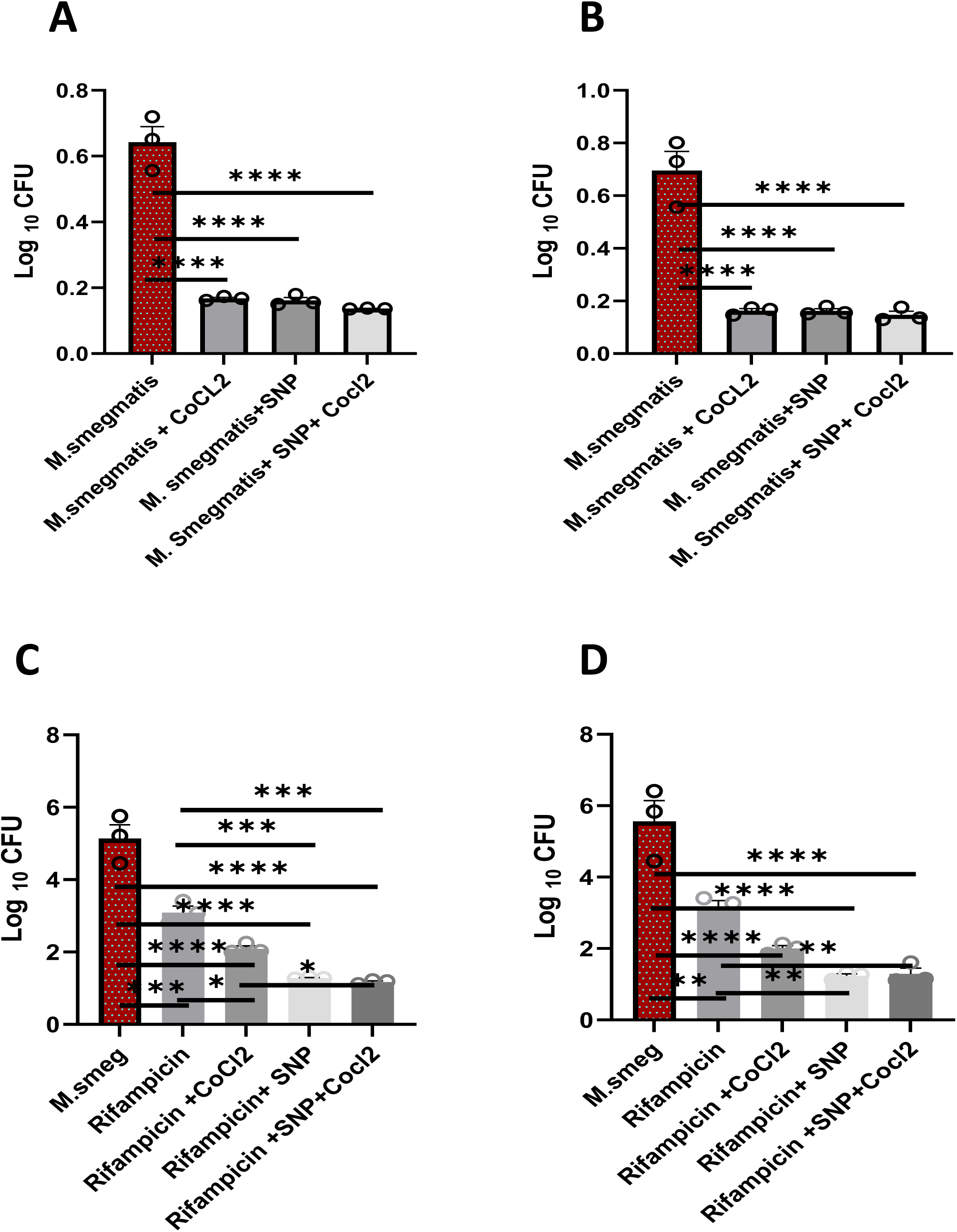
NO and Hypoxia potentially restrict mycobacterial. Mycobacteria were cultured in combination of SNP and CoCl2 and Mycobacterial growth was quantified as CFU at 48h (A) and 72h (B) post infection. Shown here the mean of CFU ± SEM quantified from 3 independent repeats at indicated time intervals. **C-D.** NO and Hypoxia influence anti tubercular action of Rifampicin. SNP and CoCL2 treated mycobacterial cultures were further treated with Rifampicin and CFU of surviving Mycobacteria were calculated at **48h (C)** and **72h (D)** respectively. Shown here the mean of CFU ± SEM quantified from 3 independent repeats at indicated time intervals

**Supplementary Figure 4.**
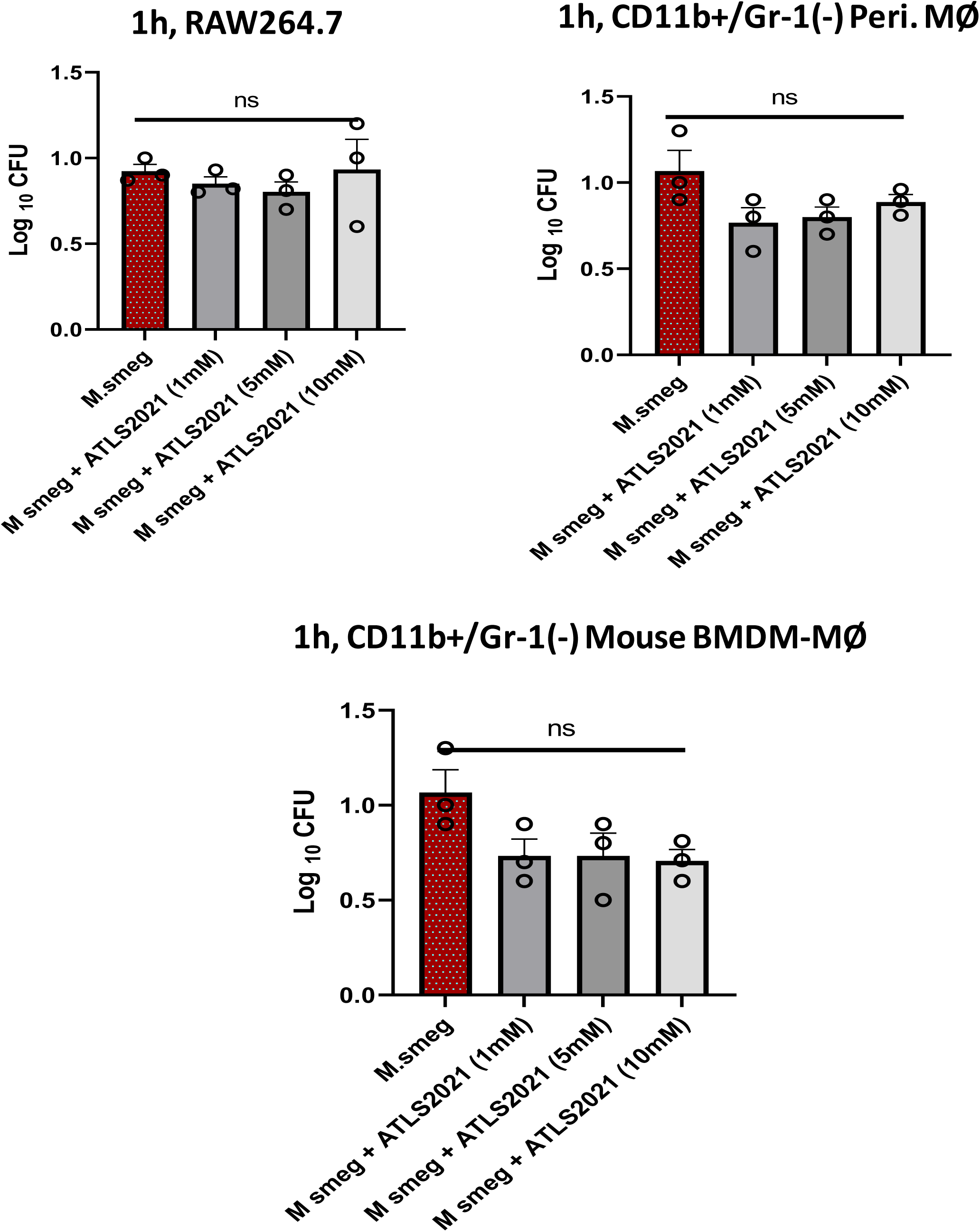
RAW macrophages were infected with M smegmatis in presence of ATLS2021 and its impact was evaluated on bacterial uptake by measuring intracellular CFU at 1h post infection. Shown here the mean of CFU ± SEM quantified from 3 independent repeats at indicated time intervals

**Supplementary Figure 5:**
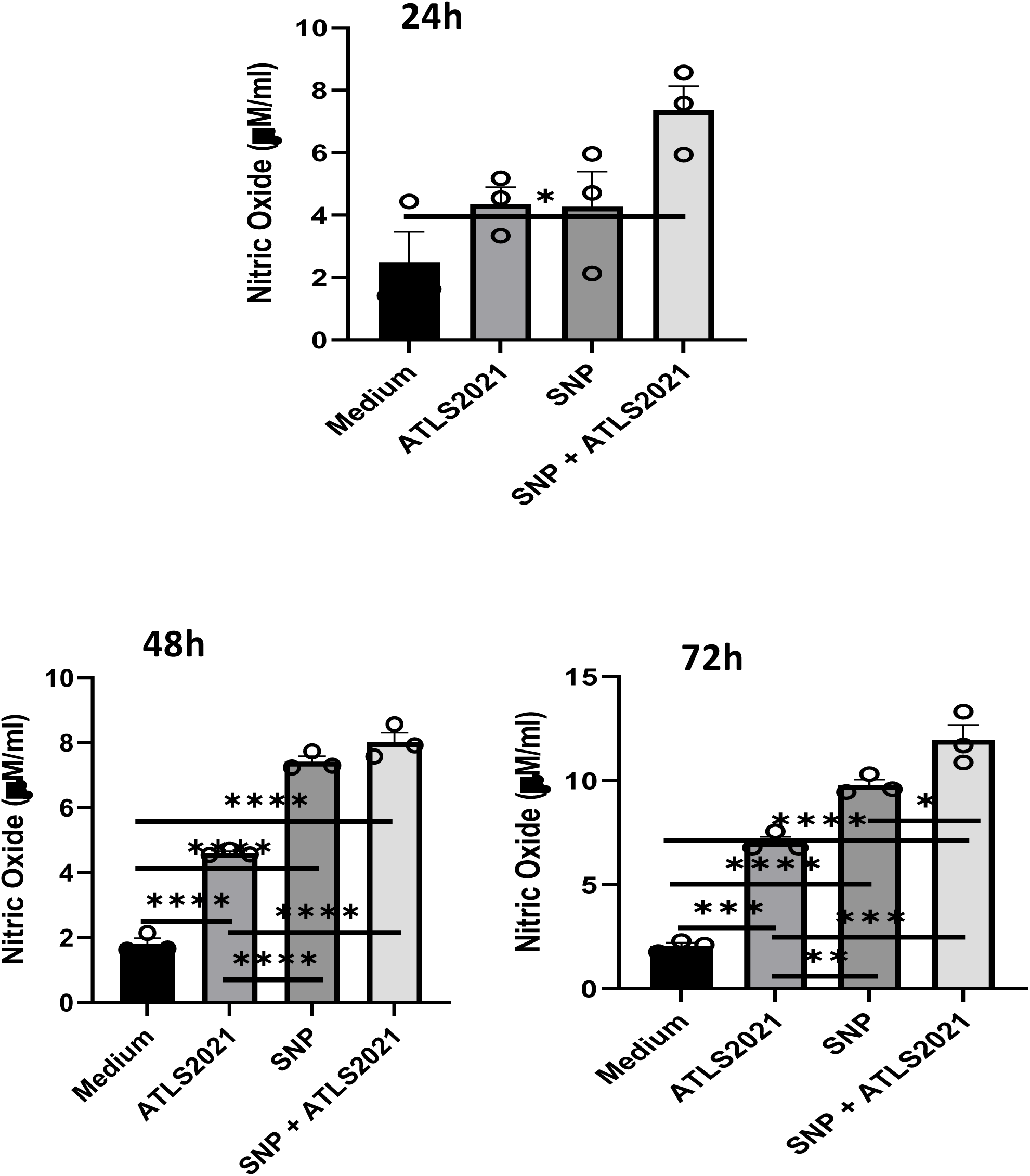
ATLS2021 enhances innate immunity. RAW macrophages were treated with ATLS2021 in presence of SNP and Nitric Oxide levels were quantified at indicated time intervals. Shown here the mean µM of NO ± SEM obtained from 3 independent repeats.

**Supplementary Figure 6.**
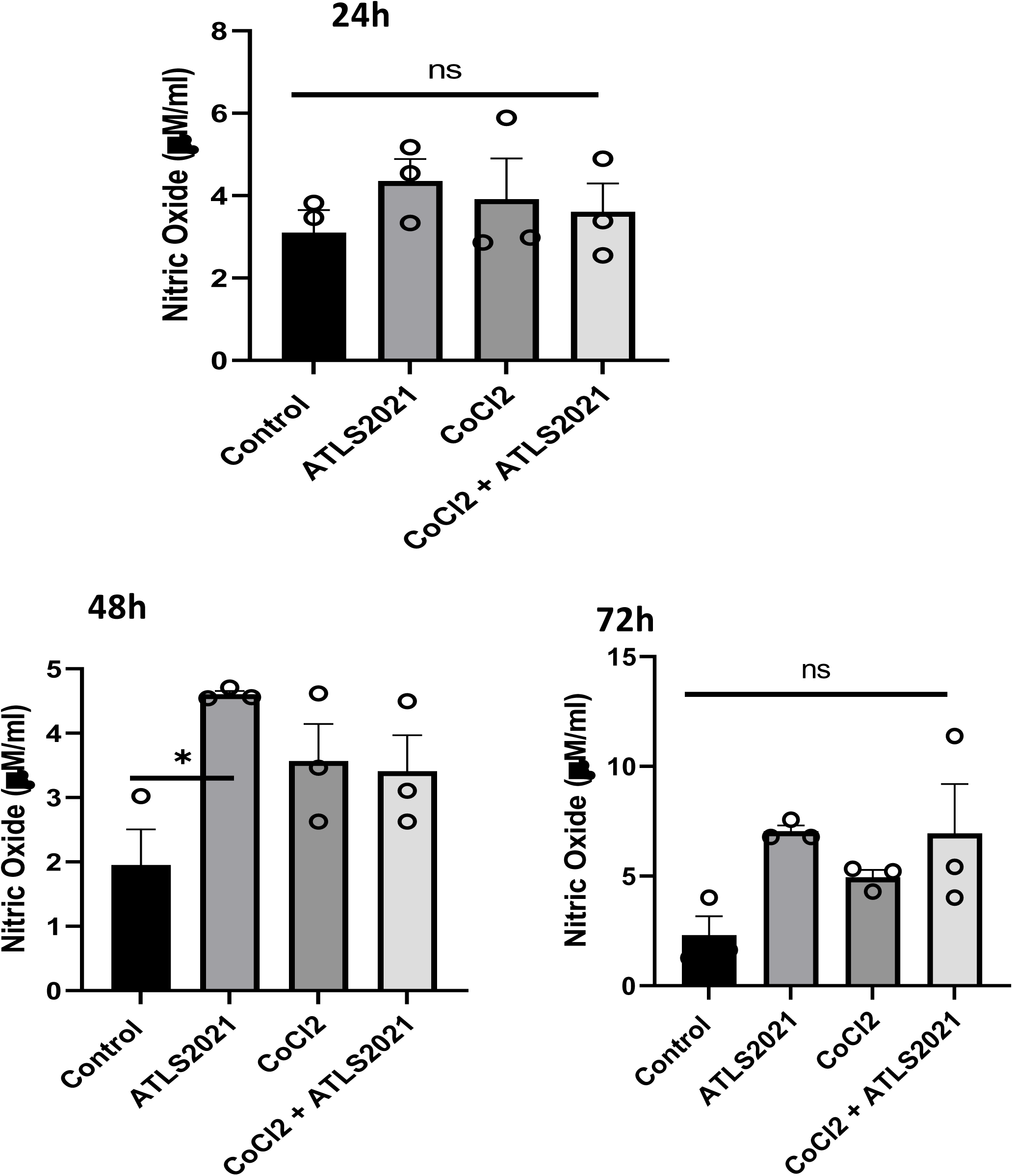
ATLS2021 enhances hypoxia induced NO in macrophages. RAW macrophages were treated with ATLS2021 in presence of COCl2 and Nitric Oxide levels were quantified at indicated time intervals. Shown here the mean µM of NO ± SEM obtained from 3 independent repeats.

**Supplementary Figure 7:**
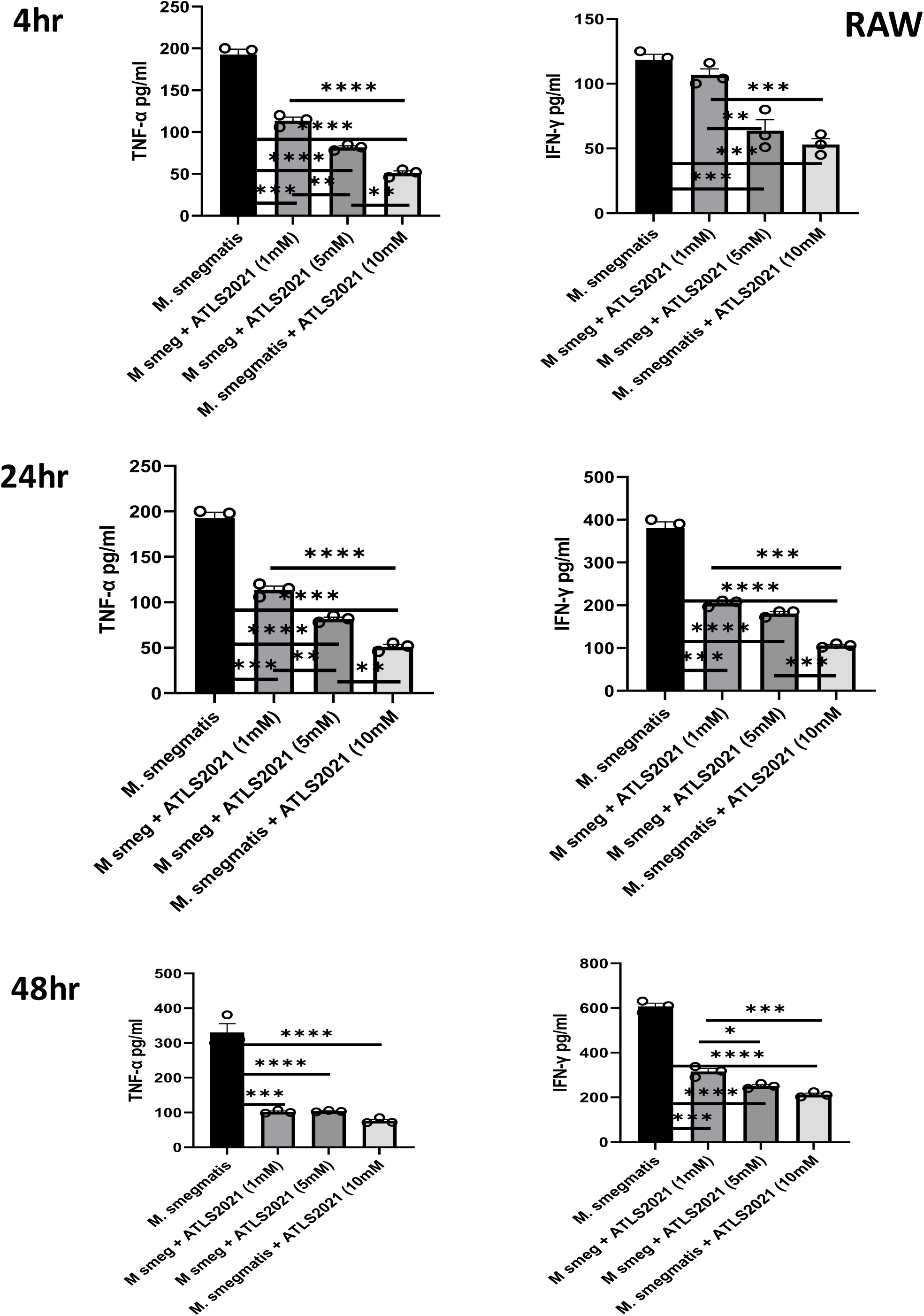
ATLS2021 normalize infection induced inflammatory stress. Culture supernatant of RAW macrophages mentioned under figure 4A were collected at indicated time point and secretion of TNF-α (left) and IFN-γ (right) levels were quantified by ELISA. Shown here the mean of *pg/ml* of indicated cytokines ± SEM quantified from three independent experiments

**Supplementary Figure 8:**
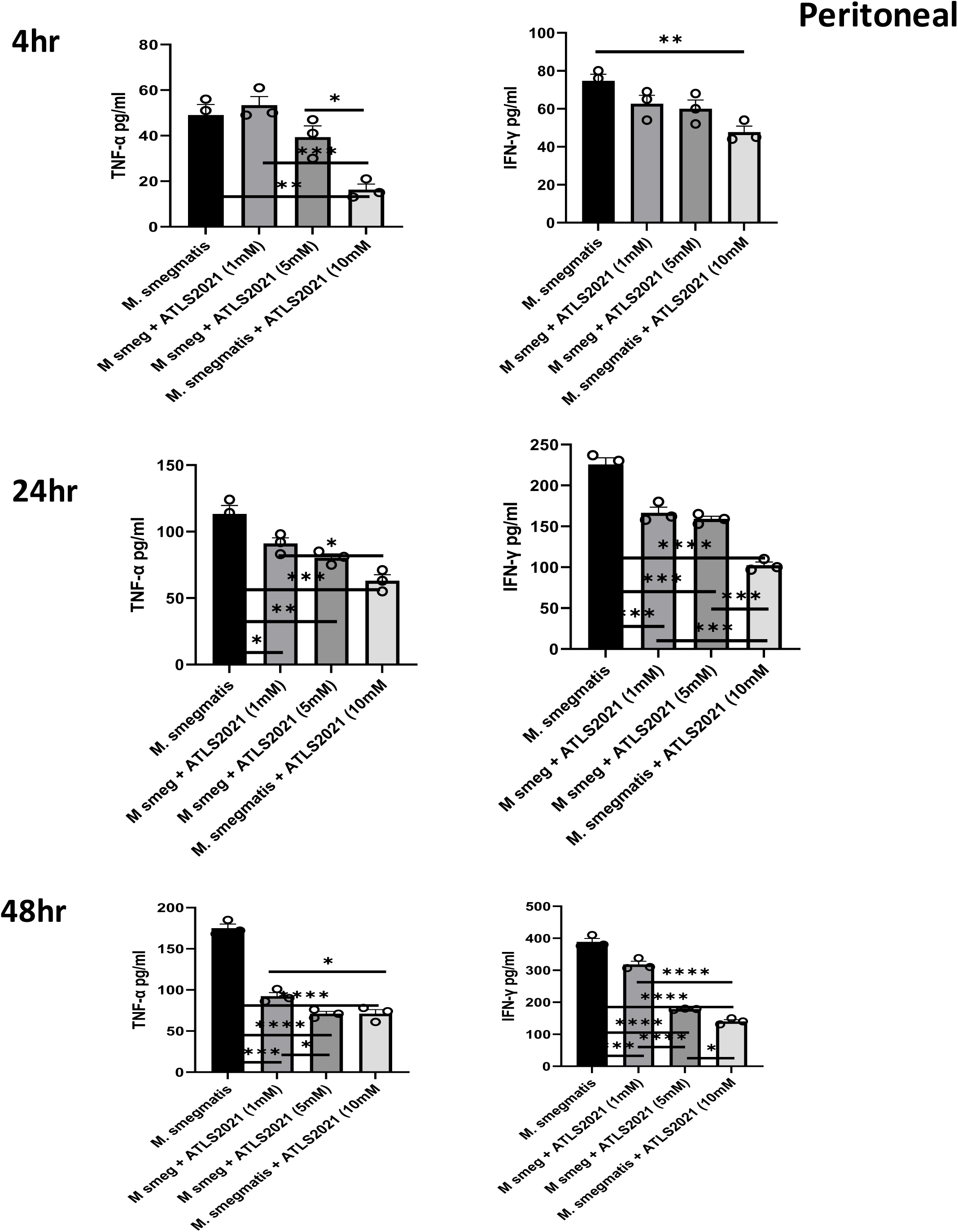
ATLS2021 normalize infection induced inflammatory stress. Culture supernatant of CD11b+/ Gr-1(−) peritoneal macrophages under figure 4B were collected at indicated time point and secretion of TNF-α (left) and IFN-γ (right) levels were quantified by ELISA. Shown here the mean of *pg/ml* of indicated cytokines ± SEM quantified from three independent experiments

**Supplementary Figure 9:**
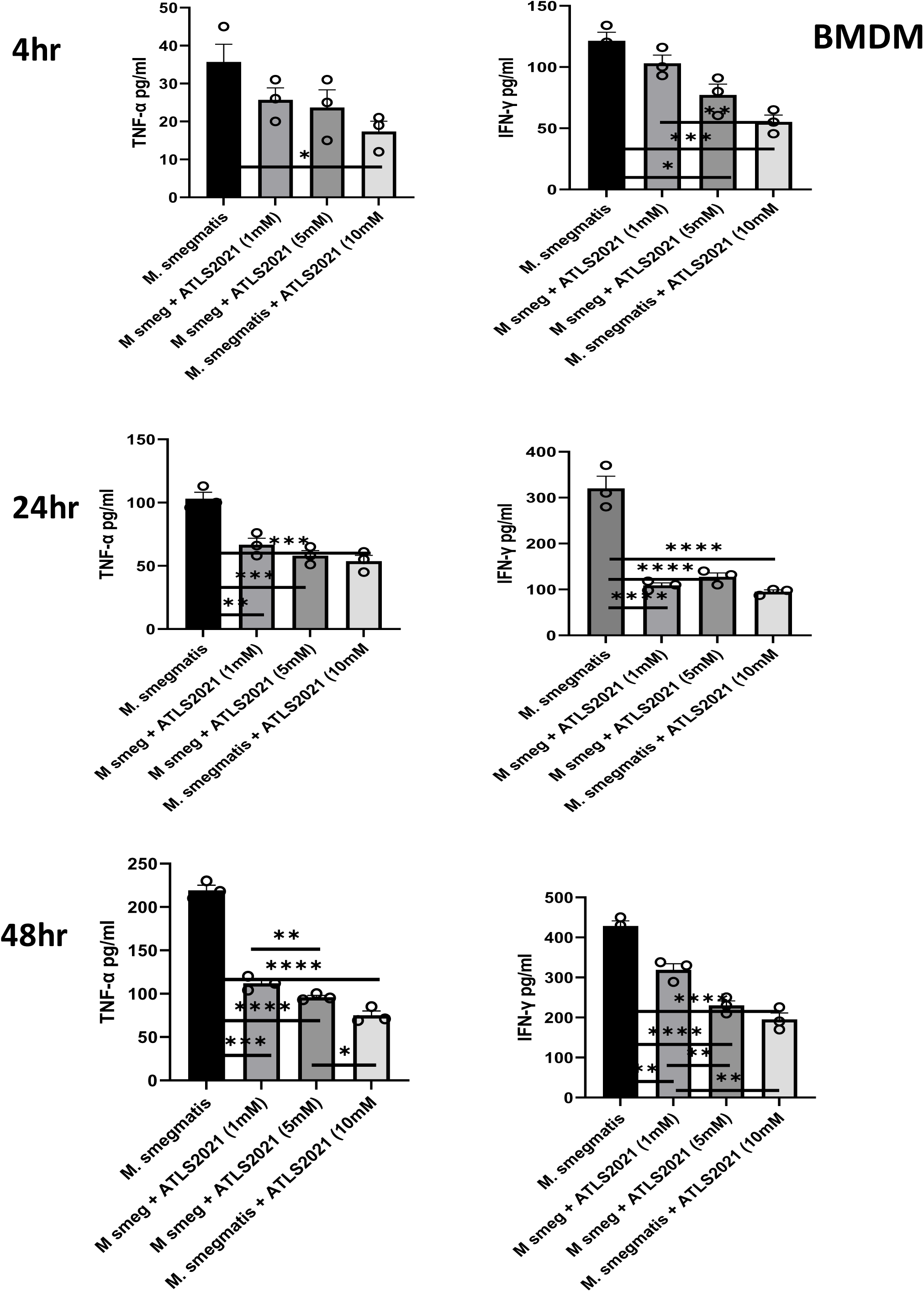
ATLS2021 normalize infection induced inflammatory stress. Culture supernatant of CD11b+/ Gr-1(−) Mouse Bone marrow derived macrophages under figure 4C were collected at indicated time point and secretion of TNF-α (left) and IFN-γ (right) levels were quantified by ELISA. Shown here the mean of *pg/ml* of indicated cytokines ± SEM quantified from three independent experiments

**Supplementary Figure 10:**
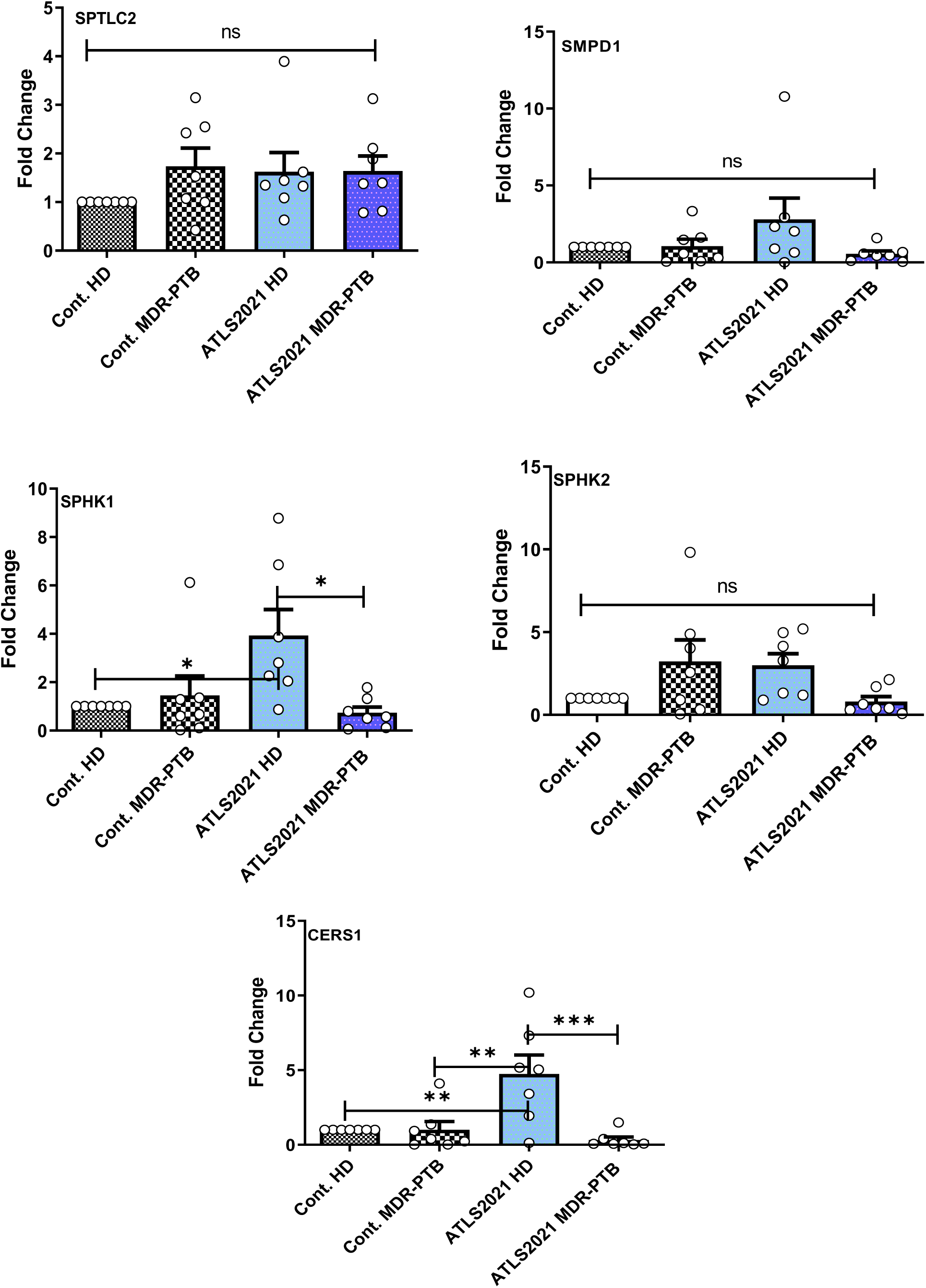
ATLS skews Sphingolipid pathways in CD14+ monocytes. CD14+ monocytes were purified from blood of Healthy and MDR-PTB patients. These cells were stimulated with ATLS2021 and gene expression of key Sphingolipid pathways components like SPTLC2, SMPD1, SMASE, SPHK1and SPHK2 were analysed by real time PCR method. Shown here the expression of indicated pathways in 6 healthy and 6 MDR PTB patients

